# Cancer cell – fibroblast crosstalk via HB-EGF/EGFR/MEK signaling promotes macrophage recruitment in squamous cell carcinoma

**DOI:** 10.1101/2023.07.13.548733

**Authors:** Giovanni Giangreco, Antonio Rullan, Yutaka Naito, Dhruva Biswas, Yun-Hsin Liu, Steven Hooper, Pablo Nenclares, Shreerang Bhide, Maggie Chon U Cheang, Probir Chakravarty, Eishu Hirata, Charles Swanton, Alan Melcher, Kevin Harrington, Erik Sahai

## Abstract

Interactions between cells in the tumor microenvironment (TME) shape cancer progression and patient outcomes. To gain new insights into how the TME influences cancer outcomes, we derive gene expression signatures indicative of signaling between stromal fibroblasts and cancer cells, and demonstrate their prognostic significance in multiple and independent squamous cell carcinoma cohorts. By leveraging information within the signatures, we discover that the HB-EGF/EGFR/MEK axis represents a hub of tumor – stroma crosstalk, promoting the expression of CSF2 and LIF and favoring the recruitment of macrophages. Together these analyses demonstrate the utility of our approach for interrogating the extent and consequences of TME crosstalk.

## Introduction

Cross-talk between cancer cells and non-malignant cells in the tumor microenvironment (TME) influences tumor growth, metastasis and therapy resistance through multiple signaling pathways and feedback mechanisms such as growth factors (TGFβ, PDGF, FGF), contact molecules (Notch, Ephrins), and inflammatory molecules (IL1, IL6, CCL12/CXCR4) ^1^. Cancer-associated fibroblasts (CAFs) promote the invasion of cancer cells, reduce the efficacy of both targeted and cytotoxic therapies and modulate immune cell recruitment and functionality ^2^. Crosstalk between cancer cells and CAFs have been demonstrated via in multiple tumors via, like oncogenic KRAS in colorectal cancer ^3^, EGFR in pancreatic ductal adenocarcinoma (PDAC) ^4^. In addition, CAFs are correlated with a pro-tumorigenic immune landscape, including higher number of tumor-promoting myeloid cells ^5^, lower numbers of tumor-infiltrating lymphocytes ^6^ and worse prognosis ^7^ ^8^. Of note, CAFs are linked to poor outcomes in squamous cell carcinoma (SCC) arising at multiple anatomical locations, including lungs (LUSC), cervix (CESC), and head and neck (HNSCC) ^8–11^. Together, these different SCC account for over 800,000 deaths per year, highlighting the need for better understanding of the disease, new therapeutic strategies, and improved tools for clinical decision making^12^.

The development of advanced sequencing techniques allows multiple inferences about the type and abundance of different TME components, including CAFs, both from bulk transcriptome and genomic methylation data ^9,13–16^. Both methods rely on the identification of cell type specific genes and the application of deconvolution strategies ultimately to infer the abundance of a particular population in a bulk dataset. However, these methods struggle to identify the functionally relevant interactions between cell types, such as signaling events ^17,18^ and the biological mechanisms associated with cell type crosstalk to be linked to patient outcomes remain incompletely understood.

Here, we propose an alternative approach to identify key players involved in tumor-stroma interaction. Instead of focusing on the abundance of CAFs or specific CAF subpopulations, we identify a signature indicative of signaling between cancer cells and CAFs. This signature is associated with worse overall survival in multiple types of SCC, pancreatic cancer, and kidney cancer. Moreover, we leverage information within the signature to identify a novel mechanism of interaction between cancer cells and CAFs. In co-culture, the RAS / MAPK pathway is strongly activated in both cell types, converging on the upregulation of Activator Protein 1 (AP-1) transcription factor (TF) components. We identify heparin-binding epidermal growth factor-like growth factor (HB-EGF) as a key mediator of cancer cell – CAF cross-talk, primarily expressed by cancer cells and able to upregulate the expression cytokines through cross-talk with CAFs. In turn, we demonstrate that this upregulation can drive attraction of macrophages, ultimately linked to worse overall survival in SCC patients (Figure S1A).

## Results

### Meta-analysis of transcriptomic data of cancer cell and cancer-associated fibroblast co-cultures identifies gene signatures with prognostic value

To identify functionally and clinically relevant gene signatures based on cancer cell – CAF cross-talk, we performed a meta-analysis of transcriptomic datasets that compare co-cultures and mono-cultures of cancer cells and CAFs. The datasets were generated under similar direct co-culture conditions using cells derived from different cancer types ^19,20^. We applied two strategies to derive gene signatures indicative of upregulated cancer cell – CAF signaling: i) selection of the most significantly enriched pathways via gene set enrichment analysis (GSEA) in co-culture for each transcriptomic dataset, followed by the selection of the up-regulated genes most frequently present in each enriched pathway (Figure 1A); ii) selection of the most up-regulated genes in co-culture for each transcriptomic dataset (Figure S1B). Using the first approach, we obtained a list of 5 genes upregulated in cancer cells and of 4 genes upregulated in CAFs upon co-culture, with one present in both. Therefore, this gene signature comprised of 8 genes (named CoCu8) (Figure 1B). The second approach led to a list of 2 genes upregulated in cancer cells and of 29 genes upregulated in CAFs upon co-culture, with one gene in common. Therefore, this gene signature consisted of 30 genes (named CoCu30) (Figure S1C).

**Figure 1:**
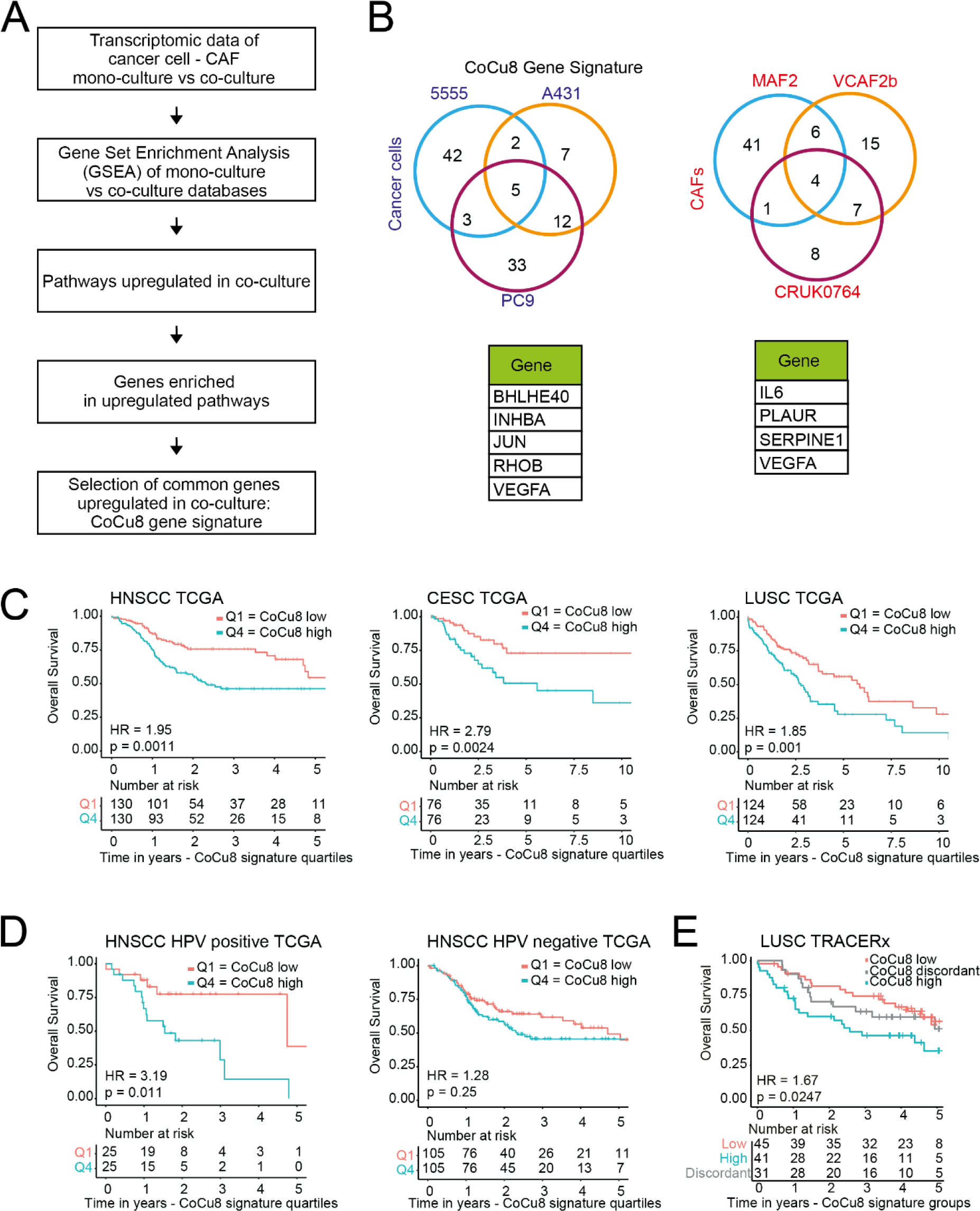
Cancer cell / CAF co-culture gene signature CoCu8 is associated with worse overall survival in multiple squamous cell carcinoma datasets. **A)** Strategy used to obtain CoCu8 gene signature. **B)** Venn diagrams of the genes upregulated in the different datasets (top) and summary table of the genes upregulated in all the datasets (bottom) for cancer cells (right) and CAFs (left). **C)** Kaplan-Meier overall survival analysis of HNSCC (right), CESC (center), LUSC (left) TCGA datasets stratified for CoCu8 first vs last quartile. Numbers at risk shown in tables below graphs. HNSCC HR=1.95 (95% Confidence Interval (CI) 1.29-2.93), p-value=0.0011. CESC HR=2.79 (95%CI 1.40-5.56), p-value=0.0024, LUSC HR=1.85 (95%CI 1.28-2.70), p-value=0.001. HR and CI were calculated using Cox regression. p-value was calculated using logRank test. **D)** Kaplan-Meier overall survival analysis of HNSCC HPV positive (right) and negative (left) TCGA datasets stratified for CoCu8 first vs last quartile. Numbers at risk shown in tables below graphs. HNSCC HPV positive HR=3.19 (95%CI 1.24-8.18), p-value=0.011. HPV negative HR=1.28 (95%CI 0.84-1.93), p-value=0.25. HR and CI were calculated using Cox regression. p-value was calculated using logRank test. **E)** Kaplan-Meier overall survival analysis of LUSC TRACERx dataset. Individual tumors stratified as high-, discordant or low-risk according to expression profile of CoCu8 signature across multiple regions, as previously described and stratified according to Biswas et al. ^60^. Briefly, patients were classified as discordant when different tumor regions from the same patient presented not unique signature levels. Below are shown the numbers at risk in years. HR=1.67 (95%CI 1.13-2.47), p-value=0.0247. HR and CI calculated using Cox regression and are referred to CoCu8 low vs CoCu8 high. p-value was calculated using logRank test.

We tested CoCu8 and CoCu30 on publicly available dataset of breast cancer – fibroblast co-cultures^21^ confirming their relevance (Figure S1D). We also tested whether CoCu8 and CoCu30 are also upregulated when cancer cells are co-cultured with other stromal cell types and for this reason, we analyzed a dataset of co-culture between 1205Lu cancer cells and HUVEC endothelial cells^22^: CoCu8 is neither enriched in cancer cells nor in endothelial cells when co-cultured, while CoCu30 shows only a weak correlation with co-culture conditions both in 1205Lu and HUVEC cells (Figure S1E). Thus, we establish new gene signatures specifically indicative of cancer cell - CAF communication.

We next sought to determine the clinical relevance of these signatures by testing the effect on patient survival for frequent cancer types in The Cancer Genome Atlas (TCGA). We found that high expression (Q4 vs Q1) of both gene signatures correlated with worse overall survival (OS) in most of the tested datasets (Figure S2A). Among them, all tested SCC datasets presented the largest effect: cervical squamous cell carcinoma (CESC, CoCu8 HR:2.79, CoCu30 HR: 2.08), HNSCC (CoCu8 Hazard Ratio (HR): 1.95, CoCu30 HR: 1.57) and lung squamous cell carcinoma (LUSC, CoCu8 HR: 1.85, CoCu30 HR: 1.78) (Figure 1C; Figure S2B), with CoCu8 consistently showing a slightly higher hazard ratio (HR) compared to CoCu30 in all three tumor types. A multivariate analysis including relevant clinical variables such as age, sex and clinical stage to evaluate the co-culture signatures effect as a continuous variable, confirmed the relevance of the signatures in these tumor types (Figure S3 for CoCu8, Figure S4 for CoCu30,). Our signature was also associated with worse survival in pancreatic and clear cell renal cell carcinoma (ccRCC) (Figure S2A). No significant link to outcome was observed in lung adenocarcinoma, breast, colorectal, bladder, or prostate cancer. Of note, given the clinical and biological differences between HPV positive and negative tumors, which warrant a different staging classification and treatment indications ^23^, we stratified HNSCC patients according to HPV status, observing that the strongest prognostic effect was visible in HPV positive samples both for CoCu8 (Figure 1D) and CoCu30 (HR: 5.47) (Figure S2C).

We validated the association between CoCu8 / CoCu30 and outcome in a second independent cohort of LUSC from the TRACERx study ^24^ (Figure 1E, Figure S2D). The multi-regional biopsies performed in the study enabled us to ask if the expression of the CoCu8 signature was uniform across tumors. Of the 117 tumors analyzed, 86 (74%) showed concordant expression of CoCu8 in all regions, which is significantly greater than would be expected based on chance (Figure 1E). Similar results were observed with CoCu30 (Figure S2D). This indicates that cancer cell-fibroblast crosstalk is typically occurring across the whole tumor. Crucially, this analysis showed that the concordant up-regulation of CoCu8 or CoCu30 across tumor regions is associated with worse prognosis. Overall, these data indicate that CoCu8 and CoCu30 signatures are associated with worse overall survival in all SCC datasets tested, therefore we decided to focus our attention on the effect of this crosstalk signature in SCC.

### The crosstalk gene signature has greater prognostic power than fibroblast abundance

Given that CoCu8 / CoCu30 reflects cancer cell-fibroblast crosstalk, the signature might be predicted to correlate with fibroblast abundance. We performed a correlation analysis of CoCu8 / CoCu30 and methyl CIBERSORT signatures in TCGA datasets, revealing that there is a statistically significant correlation between CAFs presence and CoCu8 signature (Figure S5A-B). Similar results were observed with the CoCu30 signature (Figure S5C-D). We validated these results in the TRACERx LUSC cohort and in a second independent UK_HPV positive cohort (Figure S5E-J). As methylome data was not available for these cohorts, we used fibroblast subtype gene signatures defined in a pan-cancer analysis by Galbo et al.^9^. Strong positive correlations were observed between CoCu8 / CoCu30 signature and all of the fibroblast subtypes defined both in LUSC and UK_HPV positive HNSCC (Figure S5E-J).

We speculated that our signatures of active cancer cell-fibroblast crosstalk might have better prognostic power than simply CAF abundance. To test this, we analyzed the abundance of CAFs using the methyl CIBERSORT deconvolution strategy in TCGA cohorts and probed links with overall survival^13^. This analysis indicated worse OS for HNSCC patients with higher CAF presence, but no significant differences were observed in CESC or LUSC (Figure S6A-B). Of note, both CoCu8 and CoCu30 signatures out-performed the methyl CIBERSORT method for HNSCC, CESC, and LUSC, which adds credence to our method of deriving a signature based on the interaction with fibroblasts, not simply their abundance. Overall, these data indicate that CoCu8 / CoCu30 signatures correlates with CAF abundance but have greater prognostic value than gene signatures used to infer CAF abundance.

### Pathway enrichment analysis of cancer-associated fibroblast and cancer cell co-culture reveals a consistent upregulation of AP-1 transcription factor genes

To obtain insight in the molecular basis of the cancer cell-CAF interactions, we analyzed the pathways that were enriched in all the transcriptomic datasets used to generate CoCu8 and CoCu30. This revealed upregulation of multiple pathways linked to immune regulation, stress response, and signaling (Figure 2A). Multiple genes belonging to the AP-1 transcription factor complex: *JUNB*, *FOS*, *FOSB* were strongly enriched (Figure 2B). Moreover, *PLAUR* is regulated by AP-1 factors. As our meta-analysis showed the strongest impact in the stratification of OS patients from HPV positive HNSCC, we decided to explore *JUNB*, *FOS*, *FOSB* expression levels in 4 different co-culture combinations of human HPV positive HNSCC cell lines, SCC154 and SCC47, and human oral CAFs, OCAF1 and OCAF2. Our results confirmed that these three AP-1 TFs are upregulated when cancer cells and CAFs are in direct co-culture, as compared to mono-culture (pooled RNA from both cell lines) (Figure 2C). Analysis of indirect co-cultures ^20^ separated by a 0.4μm filter indicated that CoCu8 / CoCu30 are strongly enriched with direct co-culture also when compared with indirect co-culture, implying that direct contact is required for increased AP-1 TF expression (Figure S7A).

**Figure 2:**
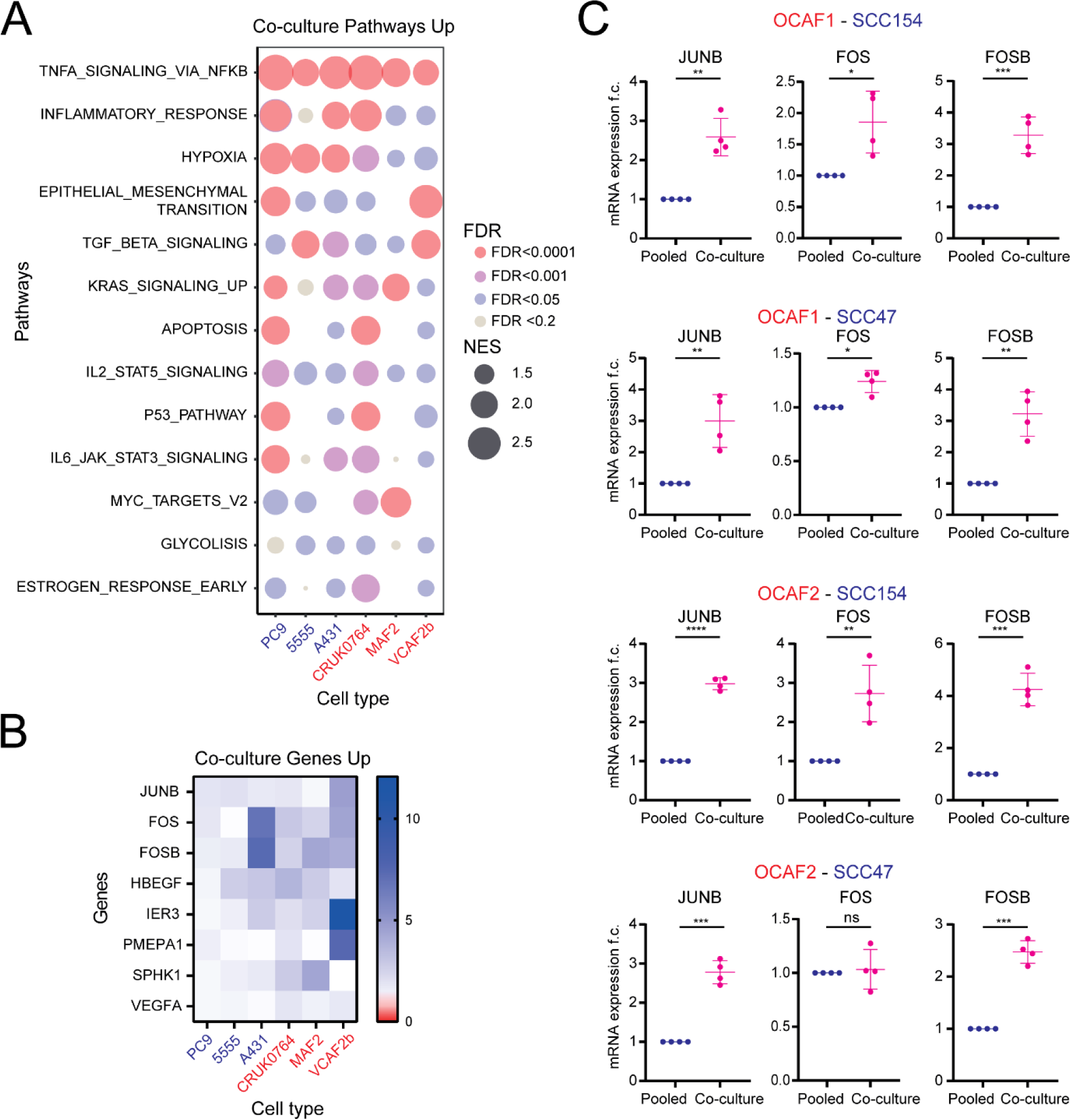
Cancer cell / CAFs co-culture upregulates AP-1 TF genes. **A)** Bubble plot of the Hallmarks pathways upregulated by cancer cells / CAFs culture. Normalized enrichment score (NES) is depicted as bubble size; False discovery rate (FDR) is depicted as color intensity. **B)** Heatmap of expression of the genes commonly upregulated upon co-culture in all the tested conditions from the TNFA_SIGNALLING_VIA_NKFB pathway. **C)** qPCR analysis of *JUNB*, *FOS* and *FOSB* genes in OCAF1/OCAF2 with SCC154/SCC47 pooled mono-cultures and co-culture after 24h. mRNA expression is reported as mean ± standard deviation (SD) fold change difference over pooled mono-culture. Genes have been normalized over the average of *GAPDH*, *ACTB* and *RPLP0* housekeeping genes. n = 4 independent experiments. Two tailed paired Student’s t-test.

### Interaction between cancer cells and cancer-associated fibroblasts is linked to increased RAS activity

We next investigated possible mechanisms underlying the upregulation of the multifunctional AP-1 TFs both in cancer cells and CAFs^25^. RAS signaling via MAPK is known to be a major driver of AP-1 gene expression^26,27^. Accordingly, we found that RAS signaling was strongly up-regulated upon direct co-culture, as indicated by the enrichment of the KRAS_SIGNALLING_UP signature (Figure 2A) and of the curated RAS84 gene signature ^28^, in all our cancer cell-CAF datasets (Figure 3A, Figure S7B).

**Figure 3:**
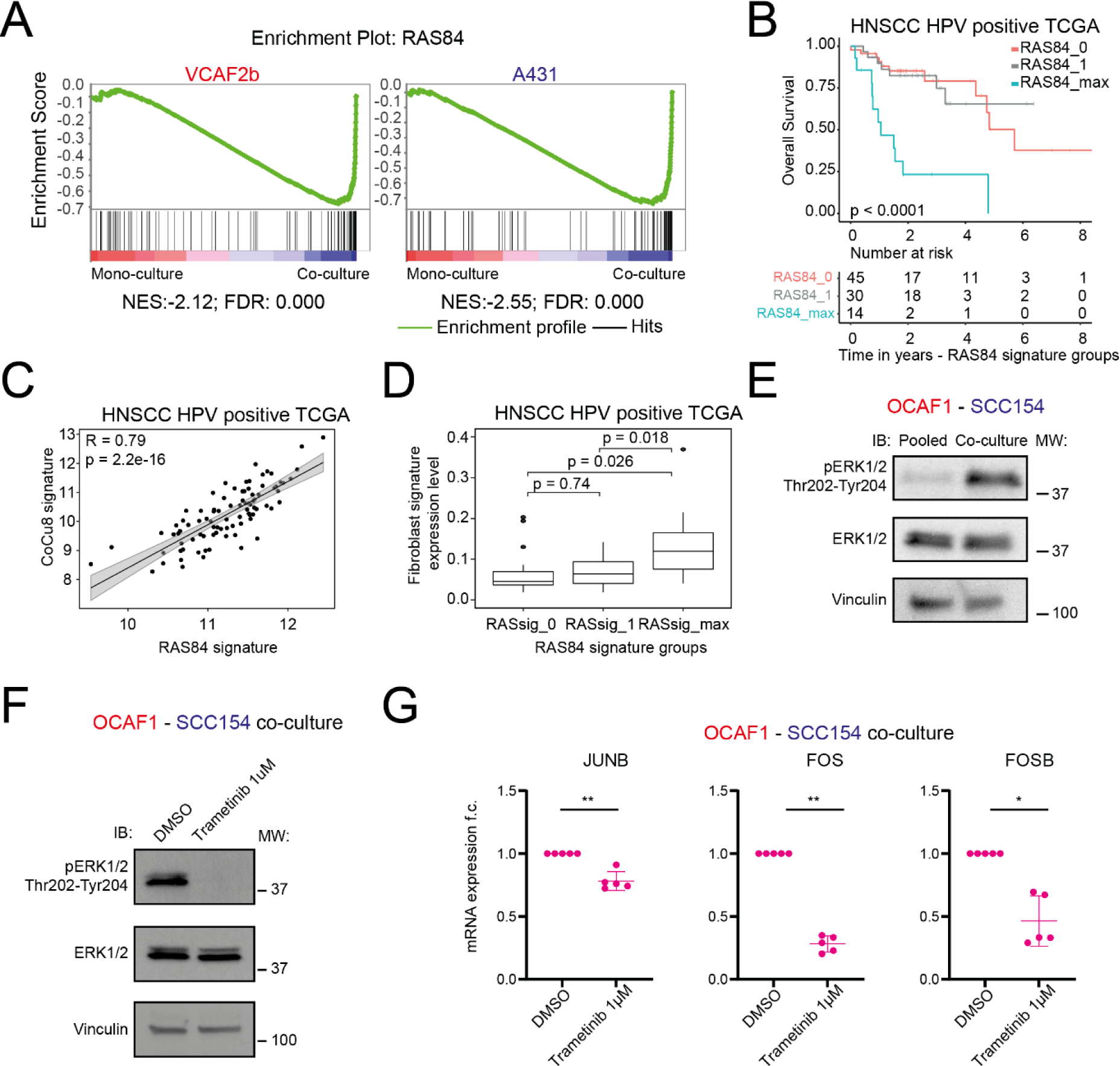
RAS activity is upregulated in cancer cell and CAFs upon co-culture. **A)** Gene set enrichment analysis (GSEA) plot of RAS84 gene signature in mono-culture and co-culture. NES and FDR are specified below each plot. **B)** Kaplan-Meier overall survival analysis of HNSCC HPV positive TCGA dataset stratified for Ras84 activity according to ^28^. Below are shown the numbers at risk in years. RAS84_0 vs RAS84_1 HR=1.01 (95%CI 0.387-2.62) p-value=0.98. RAS84_0 vs RAS84_max HR=5.85 (95%CI 2.46-13.9) p-value<0.001. HR and CI calculated using Cox regression. **C)** Correlation plot of RAS84 expression level and CoCu8 expression level in HNSCC HPV positive TCGA dataset. R is Spearman correlation coefficient. n=97. **D)** Box plot analysis of fibroblast abundance via Methyl CIBERSORT deconvolution strategy in HNSCC HPV positive TCGA dataset according to RAS84 activity. Independent Student’s t-test. **E)** Western blot analysis of OCAF1 – SCC154 pooled mono-culture vs co-culture for 48h showing the indicated antibodies. Vinculin is used as loading control. **F)** Western blot analysis of OCAF1 – SCC154 co-cultures for 48h at the indicated conditions showing the indicated antibodies. Vinculin is used as loading control. **G)** qPCR analysis of *JUNB*, *FOS* and *FOSB* genes in OCAF1 - SCC154 co-cultures for the indicated treatments after 48h. mRNA expression is reported as mean ± standard deviation (SD) fold change difference over co-culture DMSO. Genes have been normalized over the average of *GAPDH*, *ACTB* and *RPLP0* housekeeping genes. n = 4 independent experiments. Two tailed paired Student’s t-test.

To interrogate further the linkage between CAFs, the CoCu8 signature, and RAS signaling in HPV positive patients from the TCGA cohort, we stratified them according to RAS activity. We split the patient data into three groups (RAS84_0, RAS84_1 and RAS84max) according to the levels of RAS activity as performed by East et al.^28^. This analysis shows that higher RAS activity (group RAS84_max) correlates with worse OS, in a similar fashion to the effect observed with CoCu8 stratification (Figure 3B). Indeed, we observed a strong, positive correlation between RAS84 activity and CoCu8 expression in both HPV positive HNSCC cohorts (TCGA - R = 0.79, Figure 3C and UK_HPV positive cohort - R = 0.84, Figure S8A). We also observed a statistically significant enrichment in the extent of fibroblasts present when RAS activity was higher in both cohorts (Figure 3D, Figure S8B).

Activation of the RAS pathway upon cancer cell-CAF co-culture was experimentally validated by co-culturing SCC154 – OCAF1, which resulted in a strong increase in RAS-MAPK signaling, as determined by phosphorylated ERK1/2 levels (Figure 3E). To test whether the RAS-MAPK pathway is responsible for the upregulation of *JUNB*, *FOS* and *FOSB* genes, we used the MEK inhibitor, trametinib, in the SCC154-OCAF1 co-culture. We confirmed that MEK inhibition downregulates ERK1/2 activation upon co-culture (Figure 3F) and observed a significant downregulation of *JUNB*, *FOS* and *FOSB* genes (Figure 3G). Thus, multiple genes belonging to the AP-1 TF complex are upregulated when cancer cells and CAFs are co-cultured and this is mechanistically linked to the activation of RAS-MAPK kinase signaling.

### HB-EGF activation is crucial to trigger RAS pathway signaling

To explain why AP-1 TFs get upregulated in co-culture, we looked for possible activators of RAS-MAPK signaling. We noted that *HB-EGF* was among the genes upregulated in all 6 transcriptomic datasets together with *JUNB*, *FOS* and *FOSB* (Figure 2B). HB-EGF is an EGFR ligand and therefore can activate the RAS-MAPK pathway ^29,30^. We evaluated the expression levels of all seven EGFR ligands. Importantly, *HB-EGF* showed a strong and specific activation upon cancer cell - CAF co-culture (Figure 4A). Moreover, *HB-EGF* expression strongly correlates with CoCu8 in HPV positive HNSCC patients’ data from both TCGA (R=0.6, p-value=2.2e-16) and UK_HPV positive (R=0.58, p-value=1e-8) datasets (Figure 4B).

**Figure 4:**
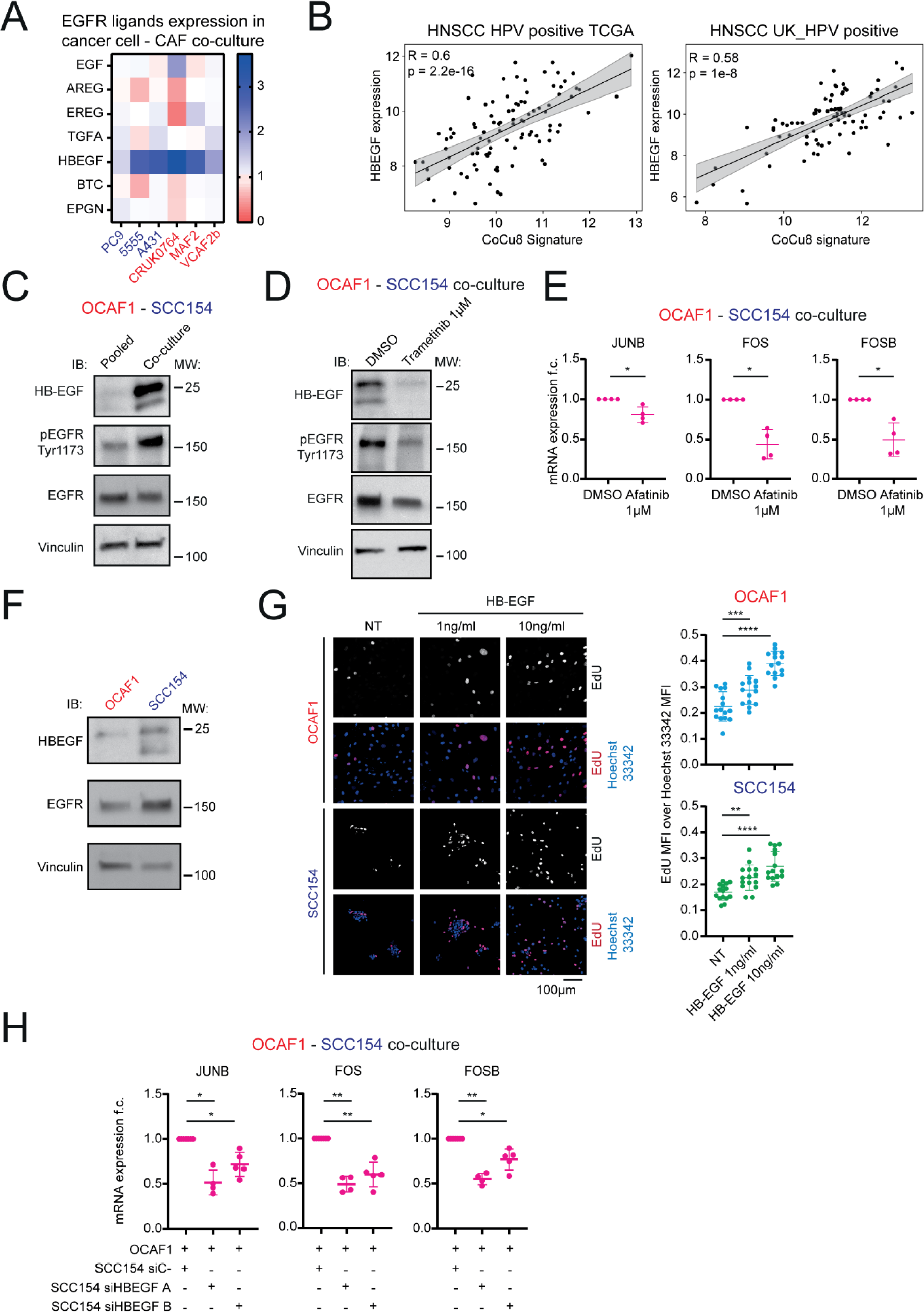
HB-EGF / EGFR axis activates AP-1 TF genes in cancer cells and CAFs upon co-culture via RAS pathway. **A)** Pattern of expression of the 7 EGFR ligands in all the tested transcriptomic datasets. **B)** Correlation plot of *HB-EGF* expression level and CoCu8 expression level in HNSCC HPV positive TCGA dataset (left, n=97) and UK_HPV positive cohort (right, n=84). R is Spearman correlation coefficient. **C)** Western blot analysis of OCAF1 – SCC154 pooled mono-culture vs co-culture for 48h showing the indicated antibodies. Vinculin is used as loading control. **D)** Western blot analysis of OCAF1 – SCC154 co-cultures for 48h at the indicated conditions showing the indicated antibodies. Vinculin is used as loading control. **E)** qPCR analysis of *JUNB*, *FOS* and *FOSB* genes in OCAF1 - SCC154 co-cultures for the indicated treatments after 48h. mRNA expression is reported as mean ± standard deviation (SD) fold change difference over co-culture DMSO. Genes have been normalized over the average of *GAPDH*, *ACTB* and *RPLP0* housekeeping genes. n = 4 independent experiments. The DMSO treated sample is the same used for Figure 3G. Two tailed paired Student’s t-test. **F)** Western blot analysis of OCAF1 and SCC154 mono-cultures after 48h showing the indicated antibodies. Vinculin is used as loading control. **G)** Proliferation assay of OCAF1 and SCC154 mono-cultures for the indicated treatments after 48h and stained with EdU and Hoechst 33342. On the left, a representative image is shown with bar graph. On the right, dot plot of mean fluorescent intensity (MFI) of EdU over Hoechst 33342 with mean ± standard deviation (SD) highlighted. Each dot is a field of view. n = 3 independent experiments. Two tailed Student’s t-test. **H)** qPCR analysis of *JUNB*, *FOS* and *FOSB* genes in OCAF1 - SCC154 co-cultures after 48h with SCC154 pre-treated with the indicated conditions. mRNA expression is reported as mean ± standard deviation (SD) fold change difference over siC-condition. Genes has been normalized over the average of *GAPDH*, *ACTB* and *RPLP0* housekeeping genes. n ≥ 4 independent experiments. Two tailed paired Student’s t-test.

HB-EGF can activate EGFR/MAPK as an un-cleaved pro-molecule at the plasma membrane ^30,31^ and, as such, signaling by membrane-bound HB-EGF could explain the need of direct cell contact to trigger the pathway. As membrane-bound HB-EGF should be expressed at about 20-25 kDa, we evaluated the cellular levels of HB-EGF in OCAF1-SCC154 co-culture and found that it was strongly upregulated at the protein level at a molecular weight previously reported as un-cleaved protein ^32^ (Figure 4C). We also observed EGFR phosphorylation was increased upon direct co-culture (Figure 4C); importantly, both these effects were abrogated by MEK inhibition (Figure 4D), suggesting the presence of a positive feedback loop involving HB-EGF / EGFR / MAPK / AP-1 upon direct co-culture. Furthermore, the EGFR inhibitor, afatinib, blocked AP-1 activation upon co-culture of SCC154-OCAF1 (Figure 4E).

These results suggest that HB-EGF might be the link that activates EGFR upon cancer cell – CAF co-culture. Therefore, we evaluated the basal expression level of EGFR and HB-EGF in SCC154 and OCAF1 mono-cultures. Interestingly, SCC154 expressed HB-EGF at much levels than OCAF1, while both cell types expressed similar levels of EGFR (Figure 4F). This suggests that both cell types can be reactive to EGF ligands, but the activation of the positive feedback loop upon direct contact requires higher levels of HB-EGF, expressed at the membrane of cancer cells. In that case, both cancer cells and CAFs should be responsive to HB-EGF treatment, albeit with potentially different downstream effects. To test this hypothesis, we incubated OCAF1 and SCC154 in mono-cultures with different concentrations of HB-EGF. Firstly, HB-EGF caused an increase in proliferation in both cell types, as shown by a higher proportion of EdU positive nuclei (Figure 4G). Moreover, SCC154 – but not OCAF1 – showed a scatter-like phenotype when treated with high doses of HB-EGF, suggestive of an epithelial to mesenchymal transition. Consistent with this, HB-EGF treatment reduced E-Cadherin staining (Figure S8C). We then tested the effect of HB-EGF treatment on known CAF markers: after treatment of OCAF1 with HB-EGF for 48h we noticed a slight but significant downregulation of *ACTA2*, a CAF and myofibroblast marker (Figure S8D). However, no effect was observed for other widely used CAF markers (*FAP*, *LRRC15*, *FN1*) (Figure S8D).

Given these data, we asked whether HB-EGF expression in cancer cells is enough to induce upregulation of AP-1 genes when cancer cells and CAFs are in co-culture: we therefore performed knock down of HB-EGF in SCC154 (Figure S8E) and then co-cultured them with OCAF1. Importantly, the downregulation of HB-EGF in SCC154 is enough to block the upregulation of *JUNB*, *FOS* and *FOSB* when cancer cells and CAFs are co-cultured (Figure 4H). These results point to an axis involving HB-EGF in cancer cells and EGFR in CAFs that activates MAPK / AP-1 inducing a positive feedback loop when cancer cells and CAFs are co-cultured.

### A paracrine HB-EGF/EGFR axis regulates cytokine expression and macrophage recruitment

To focus on the downstream effects of this crosstalk between cancer cells and CAFs and how HB-EGF could affect CAFs functions and lead to unfavorable biology, we analyzed scRNAseq dataset of HNSCC with both malignant and non-malignant samples published by Choi et al. ^33^. Interestingly, by using myofibroblast and inflammatory markers (Figure S8F), we found that EGFR is mainly expressed by inflammatory fibroblasts (iFibroblasts), but not myofibroblastic CAFs (myoFibroblasts), while HB-EGF is mainly expressed by endothelial and epithelial cells (Figure 5A).

**Figure 5:**
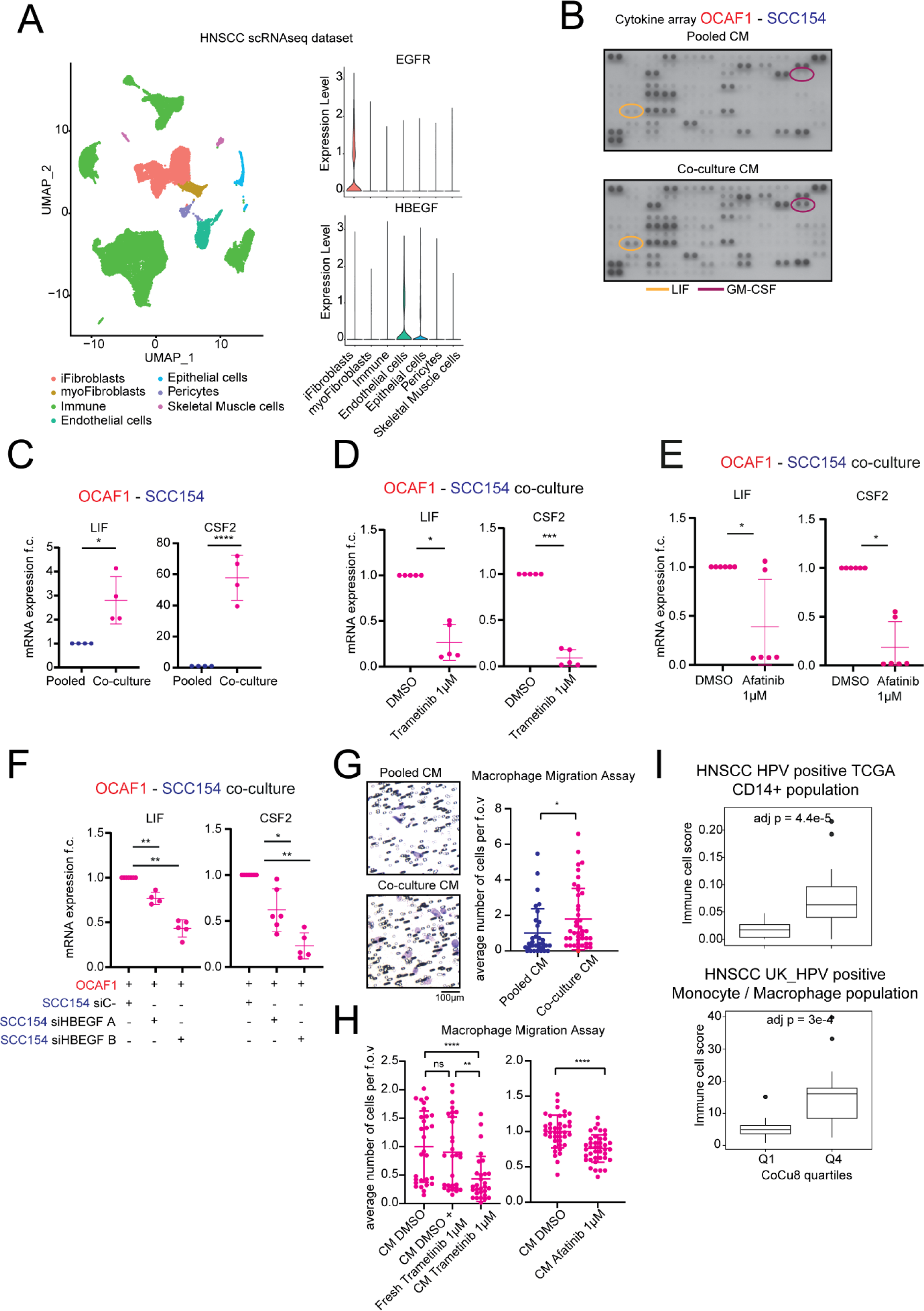
Cancer cells – CAFs co-culture induces production of specific cytokines to attract macrophages. **A)** On the right is shown scRNAseq UMAP analysis from Choi et al. ^33^. On the left is shown violin plot of EGFR and HBEGF mRNA expression levels in the indicated clusters. **B)** Cytokine array of conditioned medium from pooled mono-culture and co-culture of OCAF1 and SCC154. Highlighted relevant cytokines. n = 2 independent experiments. **C)** qPCR analysis of *LIF* and *CSF2* genes in OCAF1 – SCC154 pooled mono-culture vs co-culture after 24h. mRNA expression is reported as mean ± standard deviation (SD) fold change difference over pooled mono-culture. Genes have been normalized over the average of *GAPDH*, *ACTB* and *RPLP0* housekeeping genes. n = 4 independent experiments. Two tailed paired Student’s t-test. **D)** qPCR analysis of *LIF* and *CSF2* genes in OCAF1 - SCC154 co-cultures for the indicated treatments after 48h. mRNA expression is reported as mean ± standard deviation (SD) fold change difference over co-culture DMSO. Genes have been normalized over the average of *GAPDH*, *ACTB* and *RPLP0* housekeeping genes. n = 5 independent experiments. Paired t-test. **E)** qPCR analysis of *LIF* and *CSF2* genes in OCAF1 - SCC154 co-cultures for the indicated treatments after 48h. mRNA expression is reported as mean ± standard deviation (SD) fold change difference over co-culture DMSO. Genes have been normalized over the average of *GAPDH*, *ACTB* and *RPLP0* housekeeping genes. n = 6 independent experiments. The DMSO treated sample is the same used for Figure 5D. Two tailed paired Student’s t-test. **F)** qPCR analysis of *LIF* and *CSF2* genes in OCAF1 - SCC154 co-cultures after 48h with SCC154 pre-treated with the indicated conditions. mRNA expression is reported as mean ± standard deviation (SD) fold change difference over siC-condition. Gene has been normalized over the average of *GAPDH*, *ACTB* and *RPLP0* housekeeping genes. n ≥ 4 independent experiments. Two tailed paired Student’s t-test. **G)** Migration assay of macrophages plated in transwells with conditioned medium (CM) from OCAF1-SCC154 pooled mono-culture or co-culture. CM have been obtained after 48h culture. On the left, a representative field of view is shown with bar graph. On the right, dot plot of number of cells per field of view as mean ± standard deviation (SD). Each dot is a field of view normalized by the average of the pooled mono-culture CM sample. n = 4 different donors. Two tailed Student’s t-test. **H)** Migration assay of macrophages plated in transwells with CM from OCAF1-SCC154 co-culture with the indicated treatments. CM have been obtained after 48h culture and, for the fresh trametinib sample the drug has been added after the CM was collected. Dot plot of number of cells per field of view as mean ± standard deviation (SD) is shown. Each dot is a field of view normalized by the average of the co-culture DMSO CM sample. n = 3 different donors for trametinib effect and 4 donors for afatinib effect. Two tailed Student’s t-test. **I)** At the top, box plot analysis of CD14+ monocytic/macrophage lineage immune cell absolute score via Methyl CIBERSORT deconvolution strategy in HNSCC HPV positive TCGA dataset separate by first and last quartile of CoCu8 expression. Independent Student’s t-test, Bonferroni correction for multiple comparisons. At the bottom, box plot analysis of monocyte and macrophage immune cell score using Absolute CIBERSORT deconvolution strategy in HNSCC HPV positive UK_HPV positive dataset separate by first and last quartile of CoCu8 expression. Independent Student’s t-test.

Given that high expression of EGFR is linked to iCAFs, we performed a cytokine array of conditioned medium from pooled mono-cultures and co-cultures (Figure 5B): interestingly, we found a strong upregulation of macrophage attraction and differentiation factors (LIF and GM-CSF – gene name *CSF2* –). We validated by qPCR that *LIF* and *CSF2* are transcriptionally upregulated in co-culture (Figure 5C) and that trametinib, MEK inhibitor, treatment is enough to downregulate their expression (Figure 5D). To further investigate the involvement of EGFR in this crosstalk pathway, we performed Afatinib treatment in co-culture and observed that both *LIF* and *CSF2* are strongly downregulated by EGFR inhibition (Figure 5E). Moreover, by reducing HB-EGF expression in cancer cells and then co-culturing the cells with CAFs, we also observed strong downregulation of both *LIF* and *CSF2* expression (Figure 5F). Importantly, HB-EGF treatment in cancer cells and CAFs mono-cultures shows that: *LIF* is strongly upregulated only by CAFs, indicating that these are the cells responsible for its production when in co-culture (Figure S9A); *CSF2* was upregulated by HB-EGF treatment both in cancer cells and in CAFs (Figure S9A). In line with these, transcriptomic data of HPV positive HNSCC patients from TCGA show that there is a strong positive correlation between both *LIF* and *CSF2* mRNA and *HBEGF* mRNA expression (Figure S9B). These data establish that HB-EGF/EGFR signaling is required for the up-regulation of cytokines and that EGFR is most highly expressed in human iCAFs.

Given the established literature behind CSF2/GM-CSF and LIF involvement in macrophage biology ^34,35^, we isolated primary monocytes from peripheral blood mononuclear cells (PBMCs) from healthy donors, differentiated them into macrophages and then performed a migration assay using conditioned medium (CM) to ask if cancer cell - CAF direct CM was sufficient to increase macrophage attraction. Importantly, CM derived from the co-culture of OCAF1 and SCC154 increased the numbers of migrating macrophages, compared with pooled CM derived from each cell in monoculture (Figure 5G). We next tested if the attraction of macrophages depended on the activation of EGFR or MEK upon cancer cell-fibroblast interaction. When cancer cells and CAFs are co-cultured in the presence of trametinib, there was stark decrease in the number of migrating macrophages attracted by the CM (Figure 5H). Crucially, this was not the case when MEK inhibitor is freshly added to CM after it is harvested from the cancer cell-CAF co-culture, indicating that any residual inhibitor in the CM is not the cause of reduced macrophage attraction (Figure 5H). Moreover, blockade of EGFR using the inhibitor afatinib during the co-culture phase significantly reduced the attraction of macrophages (Figure 5H).

These data suggest that cancer cell – CAF crosstalk is sufficient to promote macrophage recruitment, therefore we asked whether CoCu8 high patients showed higher levels of macrophages. Of note, when TCGA patients’ data are separated according to CoCu8 expression, we observe a strong enrichment for cells defined by a CD14-related methylation signature (monocytes/macrophages) (Figure 5I). We found a similar pattern when patients were separated by fibroblast abundance and by RAS activity (Figure S9C). Importantly, we also observed enrichment for monocyte/macrophage lineages in our second cohort of 84 HPV positive HNSCC patients when separated for CoCu8 expression levels (Figure 5I).

To conclude, we have demonstrated that cancer cell – CAF cross-talk increases expression of different cytokines that, in turn, recruit higher numbers of macrophages. This loop is established by HB-EGF expression in cancer cells that induces a paracrine cross-talk with CAFs via EGFR dependent by RAS / MAPK activity. Activation of this pathway in both CAFs and cancer cells is needed to increase the expression of both LIF and GM-CSF. MEK inhibitor and EGFR inhibitor are sufficient to reduce the macrophage attraction.

## Discussion

The presence of CAFs in tumors correlates with worse patient survival and an immune suppressive TME in multiple tumor types ^20,36–40^, with recent studies linking different CAF subpopulations to prognosis ^41^. However, analysis based on the presence or absence of CAFs does not account for variability in the extent of functional crosstalk between cancer cells and CAFs. The approach we develop here is based on the selection of genes that are commonly upregulated in both cancer cells and CAFs upon direct cell-to-cell contact, thus focusing on the functional cancer cell-CAF interactions, rather than just on the presence of CAFs in the tumor. We applied two different strategies to select genes indicative of cancer cell-CAF interactions. The approach to define CoCu30 enriches for genes that are strongly up-regulated, which has been employed previously ^9^. To define the CoCu8 signature, we used a new approach based on the selection of a coherent set of genes linked by function. Strikingly, this method generates a signature with prognostic power in all types of SCC investigated, pancreatic, and ccRCC, with particularly strong links to outcome in HPV positive SCC. Our signature did not signify poor prognosis in breast, colorectal, or prostate cancer. We speculate that the different relevance of the signature in cancer arising in different tissues might reflect varying roles for fibroblasts in the tissue in coordinating wound healing responses, including engagement with myeloid cells.

Comparative analysis of CoCu8 and CoCu30 with annotated gene sets (KRAS SIGNALLING UP and RAS84 signature ^28^) suggested a mechanism of cross-talk between cancer cells and CAFs based on the activation of MAPK / AP-1 pathway (Figure S10). Consistent with this, the upregulation of CoCu30 genes – FOS, FOSB, JUNB, and HBEGF – required MEK activity. These data extend previous literature showing that KRAS mutation is associated with higher stromal presence ^42^ and with higher cancer cell – stromal interaction ^43^. We hypothesize that our signatures are highly prognostic in HPV positive HNSCC because it lacks oncogenic activation of EGFR or RAS, which frequently occurs in HPV negative disease ^44^. Thus, in a subset of HPV positive disease, RAS pathway activation and unfavorable downstream biology are triggered by cancer cell – fibroblast interaction.

Our analyses indicate that HB-EGF is central to the activation of MAPK signaling upon cancer cell – CAF contact. HB-EGF is the only EGF ligand to be consistently upregulated in co-culture across diverse models. Accordingly, EGFR activation is upregulated in co-culture, suggesting the presence of a positive feedback loop, including HB-EGF / EGFR / MAPK / AP-1. HB-EGF stimulation allowed us to decipher the cell type-dependent consequences of the activation of the pathway. While both cancer cells and CAFs respond to HB-EGF by activating MAPK and inducing changes in AP-1 TF expression, we observed different downstream activation mechanisms depending by the cell type. These data are consistent with the finding that AP-1 activation leads to diverse molecular and phenotypic consequences depending on the cell type studied ^45^. The modest level of HB-EGF expression by cancer cells is not sufficient to initiate these events. We propose the presence of CAFs acts as a mechanism to amplify the expression of HB-EGF, enabling a threshold for productive signaling to be exceeded. CAFs also up-regulate inflammatory cytokines more strongly than cancer cells, meaning that co-culture is required for HB-EGF to drive high levels of expression and subsequent macrophage recruitment. The mechanism through which HB-EGF is upregulated could be associated with proteolytic processing of HB-EGF at the interface between cancer cell and CAFs and will be interesting to test in future studies.

Ultimately, we link increased MAPK and EGFR activity to the chemo-attraction of macrophages. Our data provide insights into the molecular mechanism behind the correlation of CAFs and macrophages in tumors and, more generally, for links between CAFs and a pro-tumorigenic and immune-suppressive milieu ^37^. Indeed, CSF2/GM-CSF is known to be associated with macrophage enrichment and chronic inflammation and in cancer ^34^ and LIF can promote macrophage recruitment and induce a more pro-tumorigenic polarization to alter immune response during anti PD-1 therapy ^35^. Of note, CSF2/GM-CSF is produced both by cancer cells and CAFs when stimulated with HB-EGF, while LIF is specifically produced by CAFs. Distinct from our work, Mucciolo and colleagues reported that in pancreatic cancer stromal EGFR activated by AREG is involved in acquisition of pro-tumorigenic properties that favor cancer cells via myofibroblast activation^4^. This difference may reflect either difference between SCC, which is the experimental model in our work, and pancreatic cancer, or that AREG and HB-EGF may trigger different patterns of gene expression. Thus, EGFR is a critical determinant of CAF functions, with further studies required to disentangle tissue- and ligand-specific biology.

Our findings have clinical implications for patient stratification and treatment. Although HPV positive HNSCC patients typically have a better prognosis than HPV negative HNSCC patients, about 25% of these patients still have poor overall survival ^46,47^.CoCu8 / CoCu30 signatures and CAF abundance could help stratify those patients with worse prognosis within the HPV positive SCC. This improved patient stratification would be especially relevant in the context of the recent unsatisfactory efforts to de-escalate and de-intensify treatment for patients with HPV positive tumors ^48^ and could help reduce toxicity without compromising outcomes. Moreover, our results suggest that this subset of patients could benefit from a targeted approach, for example re-purposing the use of MEK or EGFR inhibitors. Indeed, trametinib – MEK inhibitor – is already used in the treatment of melanoma ^49^ and non-small-cell lung cancer ^50^ and it has been tested in phase I / II oral cavity SCC patients, showing some reduction in RAS / MAPK activity as neoadjuvant treatment ^51^. Our data argue that trametinib or EGFR inhibitors may be beneficial for HPV positive HNSCC patients with high stromal content.

In conclusion, our results demonstrate a new approach to detect biologically meaningful stromal signatures. We show that signatures based on signaling in the TME have the potential to both improve patient stratification and to identify new mechanisms of cross-talk between cancer cells and CAFs.

## Materials and methods

### Cell lines and reagents

OCAF1 and OCAF2 human fibroblasts were isolated from patient tissues of oral cancer and immortalized with lentiviral HTERT as described in ^52^. These patient samples were collected under the ethical approval REC reference 06/Q0403/125).

CRUK0764 were derived from patients with lung adenocarcinoma. These fibroblasts were established from the tumor tissue. The primary CRUK0764 was immortalized by the following infection with retroviruses expressing human telomerase reverse transcriptase.

PC9 was obtained from the Crick Institute Central Cell Services facility. PC9 were stably transfected with Lipofectamine 2000 Reagent (Thermo Fisher Scientific) according to the manufacturer’s instructions. Briefly, PC9 cell line was seeded at 50-70 % confluence in a six-well plate and transfected 2 μg of Piggybac transposase (pPBase-piggyBac) and 2 μg of mEGFP (pPBbsr2-mEGFP) plasmid DNAs. After 24h of incubation, the medium with Lipofectamine/plasmid DNA mix was replaced with a fresh medium. Cells were selected using 2 µg ml -1 blasticidin.

SCC154 (UPCI-SCC154) and SCC47 (UM-SCC47) were purchased from ATCC.

OCAF1, OCAF2, SCC154, SCC47 and CRUK0764 cells were cultured in DMEM (ThermoFisher, #41966052) containing 10% fetal bovine serum (Gibco, #10270-106), 1% penicillin/streptomycin (Invitrogen, #15140122), 1% insulin–transferrin–selenium (Invitrogen, #41400045) and kept at 37°C and 5% CO2.

PC9 were cultured in RPMI-1640 (Thermo Fisher Scientific, Rockford, IL) supplemented with 10% fetal bovine serum (Gibco, #10270-106), 1% penicillin/streptomycin (Invitrogen, #15140122) and kept at 37 °C in 5% CO2.

Cells were not allowed to reach more than 90% confluency for routine cell culture cultivation. Cell lines that are not commercially obtainable are available from the authors upon reasonable request. Routine screening for *Mycoplasma* testing was performed for all cell lines with negative results. STR profiles of human non-commercially available cell lines are included in Supplementary Table 1.

### Cell cultures conditions and treatments

Co-cultures and mono-cultures were performed with a ratio of 1:2, typically plating 5.5 x 10^5 CAFs and 2.75 x 10^5 cancer cells for a single well of a 6 well plate for the specified time point. When co-cultures were compared to pooled mono-cultures, for the mono-culture condition, same number of cells was plated but in two separated wells and then lysed together (pooled condition).

When cancer cells and CAFs mono-culture were compared among themselves, 1 x 10^6 OCAF1 and 1 x 10^6 SCC154 cells were plated in a 10cm dish.

For PC9 and CRUK0764 cells monocultures and co-cultures used for RNAseq, following 24 h co-cultures, the culture media was replaced with fresh medium with DMSO, then harvested after an additional 24 h. PC9 – CRUK0764 co-cultures were performed in a mixture of RPMI-1640 and DMEM (1:1) containing 1% fetal bovine serum (Gibco, #10270-106).

For macrophage cultivation, please see “Macrophage migration assay” section.

For cell culture treatments: drugs / factors were added when cells were plated and then added fresh after 24h. Drugs / factors used: trametinib (Selleckchem, #GSK1120212), afatinib (Selleckchem, #BIBW2992), human recombinant HB-EGF (Peprotech, #100-47).

For trametinib treatment to collect conditioned medium (CM), in order to control the effect of the drug presence regardless of its effect on secreted factors, we added fresh trametinib treatment to DMSO co-culture CM at the same concentration used for the cell co-culture treatment.

All concentrations used are specified in the figures.

RNA interference was performed with Lipofectamine RNAimax reagent from Invitrogen, according to the manufacturer’s instructions. For transient knock down of HB-EGF, cells were subjected to reverse transfection with 20 nM RNAi oligos plus forward transfection the day after, then analyzed 4 days after reverse transfection. The following RNAi oligo (Dharmacon) was used: siHB-EGF A (Cat # D-019624-02), siHB-EGF B (Cat # D-019624-03), as control the following non-targeting siRNA oligo (All Stars Negative, Quiagen, Cat # 1027281).

### Fluorescence-activated cell sorting

For PC9 – CRUK0764 RNAseq experiment, CRUK0764 were labelled with CellVue® Red Mini Kit for Membrane Labeling (Polysciences, 25567-1) according to the manufacturer’s instructions. Briefly, 1 × 10^7 cells of CRUK0764 were resuspended in the Diluent C and mixed with CellVue® Red working dye solutions (final concentration: 5 × 10^6 cells/mL, 2 × 10^6 M dye) and then incubated for 5 min at RT. Cells were washed twice with DMEM, 10% FBS medium to ensure removal of unbound fluorescence dye.

For fluorescence-activated cell sorting, cells were sorted using a flow cytometer–cell sorter BD FACSAria™ II. PC9-GFP and CRUK0764 -CellVue Red were sorted by FACS 48 h after seeding them in monoculture or direct co-culture. The cells were then trypsinised and resuspended in 3% FBS in PBS, 1 mM EDTA in preparation for sorting. Cells were separated into two populations: PC9-GFP and CRUK0764 with CellVue Red using a 488 nm laser with collection filter 530 nm/30 nm for GFP and 561 nm laser with collection filter 582 nm/20 nm for CellVue Red. Gates were designed on the basis of negative and single-color controls. All cell populations were tested for purity, and data were analyzed using FlowJo software.

### RNA sequencing analysis for co-cultures

PC9 and CRUK0764 cells were immediately centrifuged at 300 × g for 4 min to remove supernatant and add 350 µl RLT buffer (Qiagen, 79216) containing 1% β-mercaptoethanol (Sigma, M6250) and total RNA was extracted using the RNAeasy Mini kit (Qiagen, 74104; n = 3 independent experiments). Prior to library construction, the quality of total RNA was assessed by Bioanalyzer 2100 (Agilent Technologies Inc).

For RNAseq analysis: biological replicates libraries were prepared using the polyA KAPA mRNA HyperPrep Kit and sequenced on the Illumina HiSeq 4000 platform generating ∼28 million 75bp single end reads per sample. FASTQ_files were quality trimmed and adaptor removed using Trimmomatic (version 0.36) ^53^. The RSEM package (version 1.3.30) ^54^ in conjunction with the STAR alignment software (version 2.5.2a) ^55^ was used for the mapping and subsequent gene level counting of the mapped reads with respect to the Ensembl human GRCh38 (release 89) transcriptome. Normalization of raw count data was performed with the DESeq2 package (version 1. 18.1) ^56^. All the analysis was done (version 1. 18.1) ^56^ within the R programming environment (version 3. 4. 3).

To check the purity of the samples, we analyzed the resulting transcriptomic data for the expression of ‘lineage markers’. CDH1, EPCAM, CD24, and KRT genes were used as markers of carcinoma cells and for fibroblasts we used COL1A1, COL1A2, DCN, CD248, and PDGFR genes. This revealed high sample purity for all transcriptomic data, except in the PC9 – CRUK0764 experiment that had variable purity between samples. Therefore, we estimated the impurity in each sample based on the expression of the lineage marker genes and calculated the expected level of transcript if the two mono-cultures (cancer cells alone and CAFs alone) were mixed in proportion with the impurity estimate. The observed transcript in the co-culture condition was then normalized to account for the effect of contamination.

### EdU proliferation assay

The Click-iT Plus EdU Imaging Kit (Invitrogen #c10640) was used to perform the assay. Briefly, 48h after mono-cultures of OCAF1 and SCC154 were seeding, a solution with Edu 20µM was prepared and then diluted 1:1 with cell media to add EdU 10µM final concentration. After 90 minutes incubation, cells were washes twice in PBS, then fixed for 15 minutes with paraformaldehyde 3.7% and then washed twice in BSA 3%. Following this step, cells were incubated for 20 minutes with 0.5% Triton X-100 in PBS. After two BSA 3% washes, the Click-iT reaction buffer was added for 30 minutes, followed by one wash in BSA 3% and one wash in PBS. Subsequently, nuclei were stained with Hoechst 33342 at 5 µg/mL in PBS incubation for 30 minutes, followed by two PBS washes.

Samples were imaged with Zeiss 980 microscope.

### Immunofluorescence assay

The samples used to perform EdU proliferation assay have been then stained for E-Cadherin. Briefly, samples were washes twice in PBS, followed by incubation for 30 minutes in BSA 3%. Then samples were incubated over night at 4°C. After two washes in BSA 3% of 5 minutes each, samples were incubated with secondary antibody Alexa Fluor 555 in BSA 3% for 45 min. Following this step, samples were washed with PBS twice. Subsequently, nuclei were stained with Hoechst 33342 at 5 µg/mL in PBS incubation for 30 minutes, followed by two PBS washes. Samples were imaged with Zeiss 980 microscope.

### Peripheral Blood Mononuclear Cell extraction and monocytes selection

Donations of healthy blood donors were received from the Francis Crick Institute. PBMCs were isolated from whole blood using Lymphoprep (Stemcell Technologies #7811) with SepMate^TM^ density centrifugation tubes in line with manufacturer’s instructions (Stemcell Technologies #85450). Freshly isolated PBMCs were then counted before isolation of monocytes (Miltenyi Biotec #130-096-537). Monocytes were then counted for plated in normal plastic dishes.

### Cytokine array

Cytokine array used is “Proteome Profiler Human XL Cytokine Array Kit” (R&D Systems, # ARY022B) following manufacturer’s instruction. Briefly, conditioned media was isolated and filtered through a 0.4µm low protein binding PVDF Miltex syringe-driven filter (Millipore #SLHV033RS) to remove cellular debris. Media was then concentrated to 4X using Amicon® Ultra-15 and used for subsequent incubation with array.

### Macrophage migration assay

Monocytes were plated into 12-well plates (1x10^5^ cells / well) in RPMI 1640 media (ThermoFisher #12633-012) containing 10% FBS, 1% streptomycin/penicillin and 50ng/mL of M-CSF (Peprotech #300-25) and kept at 37°C and 5% CO2 for 5 days to allow macrophage differentiation. During incubation period, OCAF1 – SCC154 mono- and co-cultures were set-up for 48h. Conditioned media was isolated and filtered through a 0.4µm low protein binding PVDF Miltex syringe-driven filter (Millipore #SLHV033RS) to remove cellular debris. Media was then concentrated to 4X using Amicon® Ultra-15 centrifugal filter units (Millipore #UFC901024) and frozen into aliquots until needed. Conditioned media was added to 24-well plates, 8µm hanging cell culture inserts (Millipore #MCEP12H48) were placed on top of each well. The now differentiated macrophages were seeded inside the hanging cell culture insert and left to settle for 10 minutes before topping up media. Plates were left in the incubator for 5 hours to allow macrophages to migrate through membrane pores. After this time, the inserts were removed and the macrophages sat on top of the membrane were wiped off with a cotton bud, leaving behind the migrated macrophages at the bottom. Inserts were stained with 0.05% crystal violet for 30 minutes before washing and then imaged. Inserts were imaged using Zeiss Observer Z1 mounted with a QImaging Color camera. Quantification of crystal violet staining was carried out using ImageJ through ‘Cell Counter’ function.

### Gene Set Enrichment Analysis

Gene set enrichment analysis was performed with GSEA software v4.1.0. The dataset used to perform the comparative analysis are: RAS84 derived from ^28^, CoCu8 derived from our own analysis, Hallmarks (h.all.v7.5.symbols.gmt) for all the other analysis. All the parameters have been used as defaults except: permutation type (gene set) and metric for ranking genes (Student’s t-test). Gene signatures with a false discovery rate < 0.05 were considered as statistically significant.

### Transcriptomic data

The transcriptomic data used are: microarray data of A431 / VCAF2b under conditions of mono-cultures, co-cultures in direct contact and indirect contact are available at the Gene Expression Omnibus under record GSE121058. The microarray data of MAF2 under conditions of mono-culture and co-culture in direct contact is available at the Gene Expression Omnibus under record GSE63160. The microarray data of 5555 under conditions of mono-culture and co-culture in direct contact will be submitted at the Gene Expression Omnibus.

The RNAseq data of PC9 / CRUK0764 under conditions of mono-cultures and co-cultures in direct contact will be submitted to the European Genome-Phenome Archive before publication.

The microarray data used for HUVEC – 1205Lu analysis is available at Gene Expression Omnibus under record GSE8699.

The microarray data used for breast cancer cell lines co-culture with fibroblasts analysis is available at Gene Expression Omnibus under record GSE41678.

### Co-culture gene signature generation

CoCu8 gene signature generation: the A431/VCAF2b, 5555/MAF2, PC9/CRUK0764 co-cultures vs mono-cultures transcriptional datasets have been analyzed with GSEA (see Gene Set Enrichment Analysis method) to obtain a list of enriched pathways in co-culture with FDR < 0.05 for each condition. For each cell type, all the genes statistically upregulated have been pulled together. From this list, genes that were present in 20% or more of the enriched pathways have been selected. The results obtained for each sample have been merged according to the cell type: the three cancer cells in co-culture have been pulled together, same for the three CAFs. To select the final list, only genes present in the three different cancer cells or in the three different CAFs have been selected to generate CoCu8.

CoCu30 gene signature generation: the genes with a fold change upregulation of 1.5 or higher have been selected for each cell type upon co-culture. The results of the three cancer cells in co-culture have been pulled together, same for the three CAFs in co-culture. To select the final list, only genes present in the three different cancer cells or in the three different CAFs have been selected to generate CoCu30.

### TCGA analysis

Clinical data, RSEM (RNA-Seq by Expectation-Maximization) normalized expression data (Illumina RNASEQ platform) and Methylation data (Illumina Human Methylation 450 platform) for TCGA cohorts were downloaded from the Firebrowse website hosed by Broad Institute of MIT and Harvard. [http://firebrowse.org/]. Data downloads were all version 2016012800.0.0.

De-convolution strategies:

MethylCIBERSORT: signature matrix and mixture files were obtained using MethylCIBERSORT R package, hosted on Zenodo. The detailed origin of the signatures and the procedure to create the deconvolution strategy is explained in Chakravarthy et al. ^13^.

Absolute-CIBERSORT: To calculate the immune infiltrate per sample, the library ‘CIBERSORT’ (version 1.04 ^57^) was run within R version 3.4.3 on the RSEM normalized data and the LM22 signature using the parameters absolute=TRUE and abs_method =”no.sumto1”.

### RNA extraction and RT-qPCR

Cells were collected and lysed with RLT buffer and total RNA was extracted using the RNeasy Mini kit (Qiagen, #74104), according to the manufacturer’s protocol.

The cDNA was prepared using M-MLV reverse transcriptase (Promega, #M3682), and quantitative PCR was performed using PowerUp™ SYBR™ Green Master Mix (ThermoFisher, #A25778), using the QuantStudio 3 and 7 Real-Time PCR systems (Applied Biosystems).

Custom primers were acquired from Sigma; sequences are available in Supplementary Table 2. RNA levels were normalized using three house-keeping genes using the ΔΔC method and reported as relative fold change compared with Ctr/not treated cells/mono-culture. For each sample, technical triplicates were obtained performed and, if one of the three technical replicates was an outlier, it has been excluded. Samples with expression levels below 37 or undetected have been considered as not expressed and – in order to perform statistics – a Ct value of 40 has been assigned.

### scRNAseq analysis

scRNAseq data from Choi et al. ^33^ was downloaded from GEO (GSE181919) and analyzed using Seurat package (version 4) ^58^.

### Protein extraction, quantification and Western Blot analysis

Cells were lysed in RIPA buffer (50 mM TrisHCl, 150 mM NaCl, 1 mM EDTA, 1% Triton X-100, 1% sodium deoxycholate, 0.1% SDS), supplemented with a protease and phosphatase inhibitors (PhosSTOP tablet Roche #04906837001, cOmplete EDTA-free Roche #11873580001, 50 mM NaF). Lysis was performed directly in the cell culture plates using a cell scraper, lysates were kept for 10min on ice and then clarified by centrifugation at 16,000 g for 30 min at 4 °C.

Total protein was quantified using the bicinchoninic acid method in accordance with manufacturer’s instructions (ThermoFisher, 23225). Following protein quantification, 20 µg of sample was loaded on a 4–15% gradient Mini-PROTEAN TGX Gels (Biorad, #4561084) and transferred to a Trans-Blot Turbo Mini 0.2 µm PVDF membrane (Biorad, 1704156) for blotting. The membrane was blocked for 1 h in 5% BSA or 5% milk in TBST and then incubated overnight at 4°C or 1 h at room temperature with antibodies. The membrane was then washed before adding the horseradish peroxidase-conjugated secondary antibody (ThermoFisher), and incubating for 1 h at room temperature. The membrane was washed again before developing with Luminata Classico Western HRP substrate (Millipore, #WBLUR0100) Luminata Classico Western HRP substrate (Millipore, # WBLUF0100) and imaging. Antibody information are listed in Supplementary Table 3. All original blots are provided as source data.

### Software and visualization

Graphs were generated with Prism software (Graphpad Software v9.4.0) and R (version 4.2.1) using package ‘ggplot’ except for correlation plot in Figure S10B that was generated with cBioportal ^59^. scRNAseq data were analyzed with Seurat package (version 4).

### Statistics

Statistical analysis was performed using Prism software (Graphpad Software v9.4.0), Excel software (Microsoft Corporation v16.0) and R (version 4.2.1).

All Student’s t-tests have been performed with two tailed strategy.

P-value information: * is p-value<0.05; ** is p-value<0.01, *** is p-value<0.001, **** is p-value<0.0001.

For GSEA, we used FDR with a threshold below 0.05 to definite the significance.

Kaplan-Meier, Log-Rank and Cox regression on survival data was calculated using the R package ‘survminer’ using univariable analysis. Correlations were calculated using the Spearman method in R and the package ‘corplot’ was used to generate the graphs.

### UK_HPV positive cohort

FFPE tumor samples from 2 studies formed this cohort:

- INOVATE (MR/R015589/1 ISRCTN32335415), a prospective sample collection study in patients with T1-T2/N1-3 or T3-T4/N0-3 oropharyngeal cancer (AJCC TNM classification 7.0) receiving treatment with radical radiotherapy with or without additional platin-based chemotherapy.
- INSIGHT-2 (C7224/A23275 NCT04242459), a prospective study of optimizing radiation therapy in head and neck cancers using functional image-guided radiotherapy and novel biomarkers. INOVATE was approved by the London -Bloomsbury Research Ethics Committee (19/LO/1558) and INSIGHT-2 was approved by London – Queen Square Research Ethics Committee (19/LO/0638).

Written informed consent was obtained from all participants prior to any study procedure.

### UK_HPV positive cohort RNAseq and data analysis

Baseline diagnostic biopsies embedded in paraffin blocks were obtained from the above-mentioned cohort. Relevant tumor sections were selected and RNA was extracted from 3-4 slides using the Qiagen AllPrep® DNA/RNA FFPE kit (#80234). Ribosomal RNA was depleted using QIAGEN FastSelect rRNA H/M/R kit (#334375). RNA sequencing libraries were prepared using the NEBNext Ultra II RNA Library Prep Kit (#E770) for Illumina following manufacturer’s instructions. The sequencing libraries were multiplexed and loaded on the flow cell on the Illumina NovaSeq 6000 instrument according to manufacturer’s instructions. The samples were sequenced using a 2x150 Pair-End (PE) configuration v1.5 for an estimated output of ∼50M paired-end reads per sample. Image analysis and base calling were conducted by the NovaSeq Control Software v1.7 on the NovaSeq instrument. Raw sequence data (.bcl files) generated from Illumina NovaSeq was converted into fastq files and de-multiplexed using Illumina bcl2fastq program version 2.20. One mismatch was allowed for index sequence identification. Sample adequacy was confirmed using FASTQC, low quality bases and reads were trimmed using Trimmomatic, we run Hisat2-Stringtie for alignment.

RNAseq was performed on 103 patient samples, of which 7 were from the INSIGHT2 and 96 from INOVATE. The data from RNAseq was analyzed to identify samples with presence of HPV (by aligning the unmapped sequences to the whole HPV16 genome sequence obtained from GEO using HISAT2 and StingTie) and these samples were classed as HPV positive. 84 samples (77 INOVATE and 7 INSIGHT2) were classified at HPV positive and RNAseq data from these was used for analysis in this study.

### TRACERx cohort

Tumor samples used in this study were collected from LUSC patients enrolled as a part of TRACERx study (accession code: NCT01888601) which is sponsored by University College London (UCL/12/0279) and has been approved by an independent research ethics committee (13/LO/1546). Multiple regions were sampled per tumor and processed as described by Frankell et al. ^24^ yielding whole-RNA sequencing data for 295 regions from 117 LUSC patients. Expression count and transcript per million (TPM) were quantified by the RSEM package ^54^. Genes with expression level of at least 1 TPM in at least 20% of the samples were included. A variance stabilizing transformation (VST) was then applied to filtered count using the DESeq2 package ^56^.

## Supplementary tables

Supplementary Tables 1, 2 and 3 are provided with this article.

## Data accessibility

The transcriptomic data for UK_HPV positive cohort is part of ongoing clinical trials, therefore the data cannot be deposited in a public repository until the trial is finalised. Data can be shared upon reasonable request following corresponding Ethical Research Committee approval following the ICR-Clinical Trials and Statistics Unit policy.

## Authors contributions

G.G. and A.R. conceptualized and designed the research, with guidance from E.S. GG performed the experimental analysis and identified the cancer cell – CAF crosstalk gene signature strategy. A.R. performed the clinical data analysis on TCGA and UK_HPV positive cohorts and performed the deconvolution strategies. G.G., A.R., S.H., E.H., Y.N., P.N. and P.C. acquired and analyzed the data, except for TRACERx data that were analyzed by D.B. and UH.L.. C.S., A.M., K.H., S.B., M.C. and E.S. provided funding and access to samples and reagents. G.G., A.R. and E.S. wrote the manuscript. All authors reviewed and edited the manuscript. All authors authorized the final version.

## Declaration of Interests and acknowledgments

We thank the Crick healthy donors’ program and donators for blood donations from the Francis Crick Institute. We thank Sophie de Carne and Phil East for useful discussions around RAS signature and RAS signaling.

G.G. is funded by Merck Sharp & Dohme Corp, New Jersey, USA (LKR190557). A.R. acknowledges funding from the Spanish Society for Medical Oncology (Beca Fundación SEOM), a CRUK accelerator grant to The Francis Crick Institute, and the Royal Marsden NIHR/BRC Bridge funding program. E.H. is supported by the Japan Society for the Promotion of Science (JSPS) Kakenhi Grant [20H03510]. Y.N was supported by a Grant-in-aid for JSPS Overseas Research Fellowship (No. 201860634).

K.H. and A.M. acknowledge funding by the Wellcome Trust, ICR/RM NIHR Biomedical Research Centre, The Institute of Cancer Research/Royal Marsden Hospital Centre for Translational Immunotherapy, CRUK Head and Neck Programme Grant (C7224/A23275) and ICR/RM CRUK RadNet Centre of Excellence (C7224/A28724). Disclosures: Honoraria: Arch Oncology (Inst), AstraZeneca (Inst), BMS (Inst), Boehringer Ingelheim (Inst), Codiak Biosciences (Inst), F-Star Therapeutics (Inst), Inzen Therapeutics (Inst), Merck Serono (Inst), MSD (Inst), Oncolys Biopharma (Inst), Pfizer (Inst), Replimune (Inst), VacV Biotherapeutics (Inst); Consulting or Advisory Role: Arch Oncology (Inst), AstraZeneca (Inst), BMS (Inst), Boehringer Ingelheim (Inst), Inzen Therapeutics (Inst), Merck Serono (Inst), MSD (Inst), Oncolys BioPharma (Inst), Replimune (Inst); Speakers’ Bureau: BMS (Inst), Merck Serono (Inst), MSD (Inst); Research Funding: AstraZeneca (Inst), Boehringer Ingelheim (Inst), Merck Sharp & Dohme (Inst), Replimune (Inst).

S.B. declares no conflict of Interest and funding support from ICR/RM NIHR Biomedical Research Centre, CRUK Head and Neck Programme Grant (C7224/A23275), ICR/RM CRUK RadNet Centre of Excellence (C7224/A28724) and the Medical Research Council Developmental Pathway Funding Scheme [MR/R015589/1].

D.B. was supported by funding from a Cancer Research UK (CRUK) Early Detection and Diagnosis Project award, the Idea to Innovation (i2i) Crick translation scheme supported by the Medical Research Council, the National Institute for Health Research Biomedical Research Centre and the Breast Cancer Research Foundation (BCRF). D.B. reports personal fees from NanoString and AstraZeneca, and has a patent PCT/GB2020/050221 issued on methods for cancer prognostication.

C.S. acknowledges grant support from AstraZeneca, Boehringer-Ingelheim, Bristol Myers Squibb, Pfizer, Roche-Ventana, Invitae (previously Archer Dx Inc - collaboration in minimal residual disease sequencing technologies), and Ono Pharmaceutical. He is an AstraZeneca Advisory Board member and Chief Investigator for the AZ MeRmaiD 1 and 2 clinical trials and is also Co-Chief Investigator of the NHS Galleri trial funded by GRAIL and a paid member of GRAIL’s Scientific Advisory Board. He receives consultant fees from Achilles Therapeutics (also SAB member), Bicycle Therapeutics (also a SAB member), Genentech, Medicxi, Roche Innovation Centre – Shanghai, Metabomed (until July 2022), and the Sarah Cannon Research Institute. C.S has received honoraria from Amgen, AstraZeneca, Pfizer, Novartis, GlaxoSmithKline, MSD, Bristol Myers Squibb, Illumina, and Roche-Ventana. C.S. had stock options in Apogen Biotechnologies and GRAIL until June 2021, and currently has stock options in Epic Bioscience, Bicycle Therapeutics, and has stock options and is co-founder of Achilles Therapeutics.

C.S. holds patents relating to assay technology to detect tumor recurrence (PCT/GB2017/053289); to targeting neoantigens (PCT/EP2016/059401), identifying patent response to immune checkpoint blockade (PCT/EP2016/071471), determining HLA LOH (PCT/GB2018/052004), predicting survival rates of patients with cancer (PCT/GB2020/050221), identifying patients who respond to cancer treatment (PCT/GB2018/051912), US patent relating to detecting tumor mutations (PCT/US2017/28013), methods for lung cancer detection (US20190106751A1) and both a European and US patent related to identifying insertion/deletion mutation targets (PCT/GB2018/051892). C.S. is a Royal Society Napier Research Professor (RSRP\R\210001). This work was supported by the Francis Crick Institute that receives its core funding from Cancer Research UK (CC2041), the UK Medical Research Council (CC2041), and the Wellcome Trust (CC2041). For the purpose of Open Access, the author has applied a CC BY public copyright licence to any Author Accepted Manuscript version arising from this submission. C.S. is funded by Cancer Research UK (TRACERx (C11496/A17786), PEACE (C416/A21999) and CRUK Cancer Immunotherapy Catalyst Network); Cancer Research UK Lung Cancer Centre of Excellence (C11496/A30025); the Rosetrees Trust, Butterfield and Stoneygate Trusts; NovoNordisk Foundation (ID16584); Royal Society Professorship Enhancement Award (RP/EA/180007); National Institute for Health Research (NIHR) University College London Hospitals Biomedical Research Centre; the Cancer Research UK-University College London Centre; Experimental Cancer Medicine Centre; the Breast Cancer Research Foundation (US) BCRF-22-157; Cancer Research UK Early Detection an Diagnosis Primer Award (Grant EDDPMA-Nov21/100034); and The Mark Foundation for Cancer Research Aspire Award (Grant 21-029-ASP). This work was supported by a Stand Up To Cancer-LUNGevity-American Lung Association Lung Cancer Interception Dream Team Translational Research Grant (Grant Number: SU2C-AACR-DT23-17 to S.M. Dubinett and A.E. Spira). Stand Up To Cancer is a division of the Entertainment Industry Foundation. Research grants are administered by the American Association for Cancer Research, the Scientific Partner of SU2C. C.S. is in receipt of an ERC Advanced Grant (PROTEUS) from the European Research Council under the European Union’s Horizon 2020 research and innovation programme (grant agreement no. 835297).

E.S. is supported by the Francis Crick Institute which receives its core funding from Cancer Research UK (CC2040), the UK Medical Research Council (CC2040), and the Wellcome Trust (CC2040). E.S. is additionally funded by the European Research Council ERCAdG CAN_ORGANISE 101019366. E.S. declares research funding from Merck Sharp & Dohme and Astrazeneca and consultancy work for Phenomic AI and Theolytics. For the purpose of Open Access, the author has applied a CC BY public copyright license to any Author Accepted Manuscript version arising from this submission.

RNA sequencing of the UK_HPV positive cohort was funded by the NIHR through the RMH BRC Pump Priming theme (grant B140).

The INOVATE study was funded by the Medical Research Council Developmental Pathway Funding Scheme [MR/R015589/1]. The INSIGHT-2 trial was supported by CRUK Head and Neck Programme Grant (C7224/A23275).

This study represents independent research supported by the National Institute for Health Research (NIHR) Biomedical Research Centre at the Royal Marsden National Health Service Foundation Trust and the Institute of Cancer Research, London, UK.

## Abbreviations

Activator protein 1: AP-1
Cancer associated fibroblasts: CAF
Cervical squamous cell carcinoma: CESC
Clear cell Renal Cell Carcinoma: ccRCC
Chemokine (C-C) motif ligand: CCL
Confidence Interval: CI
Conditioned medium: CM
False discovery rate: FDR
Fibroblast growth factor: FGF
Gene-set enrichment analysis: GSEA
Hazard ratio: HR
Head and neck squamous cell carcinoma: HNSCC
Heparin-binding epidermal growth factor-like growth factor: HB-EGF
Human papillomavirus: HPV
Interleukin: IL
Lung squamous cell carcinoma: LUSC
Overall Survival: OS
Mean fluorescent intensity: MFI
Normalized enrichment score: NES
Non-treated: NT
Pancreatic ductal adenocarcinoma: PDAC
Peripheral blood mononuclear cell: PBMC
Platelet derived growth factor: PDGF
Single-cell RNA sequencing: scRNAseq
Standard deviation: SD
Short tandem repeats: STR
Squamous cell carcinoma: SCC
The Cancer Genome Atlas: TCGA
Transcription factor: TF
Transforming Growth Factor: TGF
Tumor microenvironment: TME
Tumor necrosis factor: TNF

---

**Figure S1:**
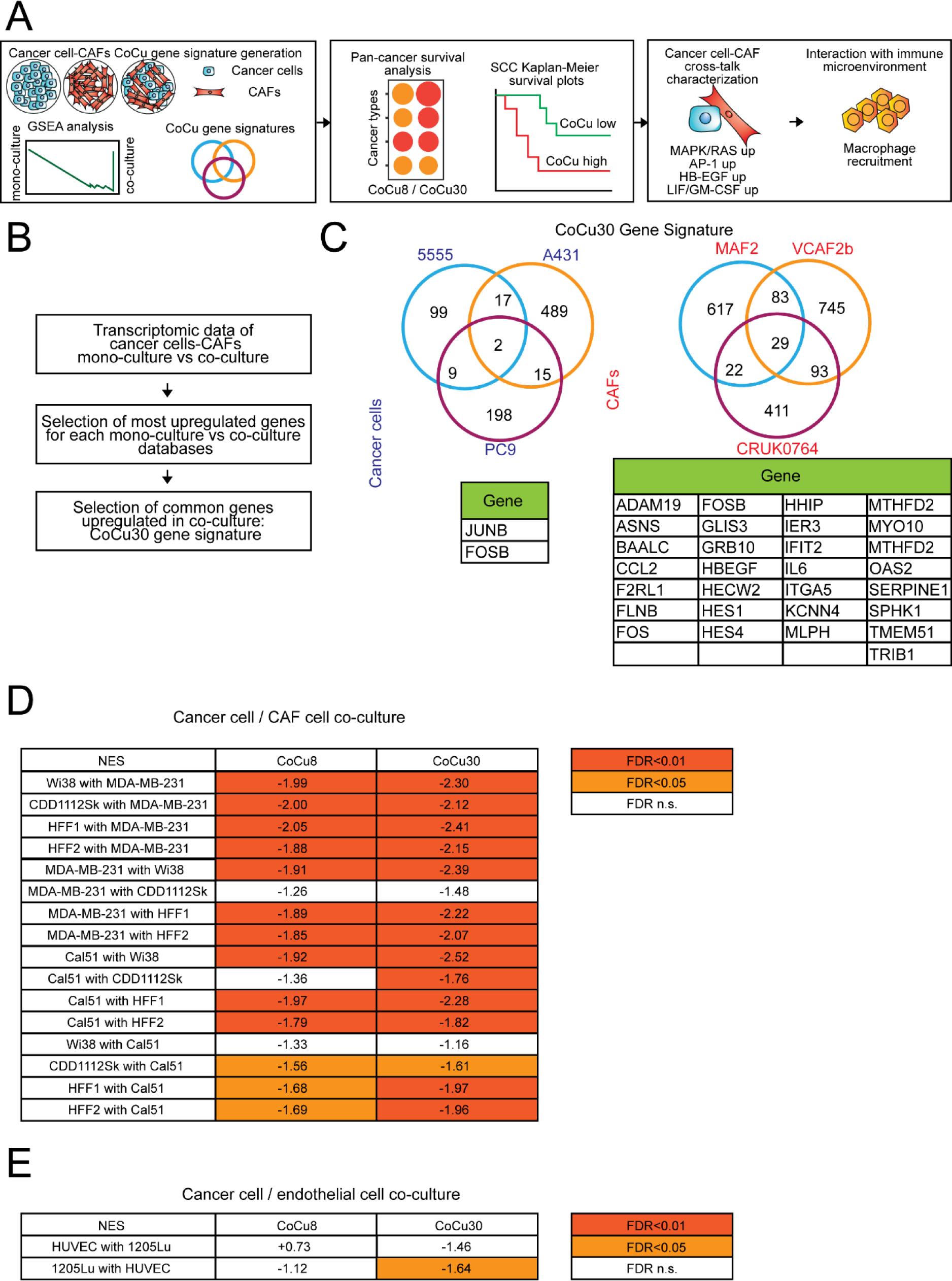
Generation and validation of CoCu8 / CoCu30 gene signatures. **A)** Flow chart description of the manuscript is provided. **B)** Strategy used to obtain CoCu30 gene signature. **C)** Venn diagram of the genes upregulated in the different datasets (top) and a table to summarize the genes constantly upregulated in the datasets (bottom) for cancer cells (left) and CAFs (right). **D)** Table with Normalized Enrichment Score (NES) values of different combinations of cancer cells and CAFs breast cancer cell lines from Rajaram et al. ^21^. Negative values represent enrichment towards co-culture condition. Color legend is shown. **E)** Table with NES values of CoCu8 and CoCu30 gene signatures for cancer cells and endothelial cells mono-culture vs co-culture available from Stine et al. ^22^. Negative values represent enrichment towards co-culture condition. Color legend is shown.

**Figure S2:**
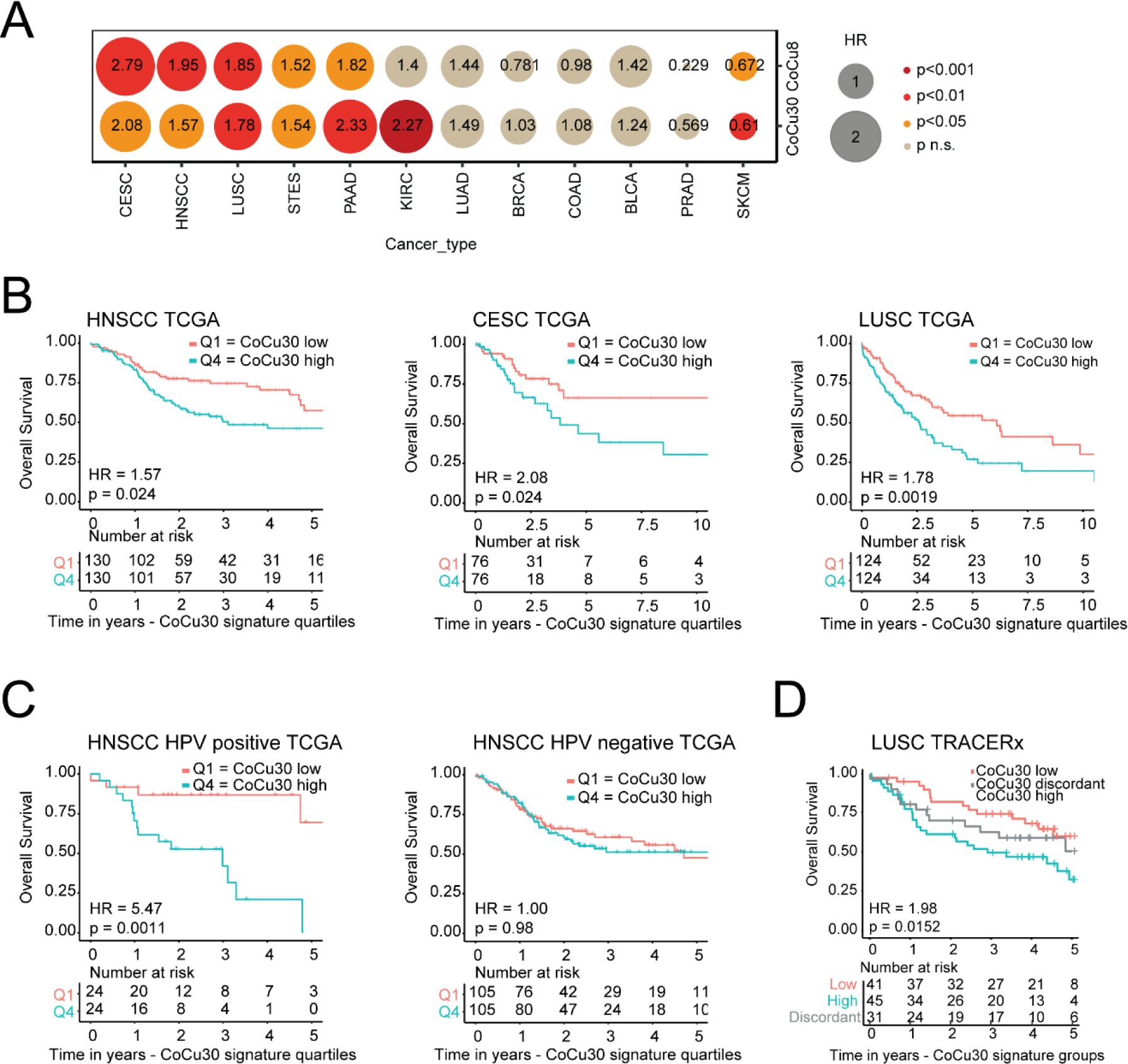
CoCu8 / CoCu30 gene signatures are associated with worse overall survival in multiple squamous cell carcinoma datasets. **A)** Bubble plot hazard ratios and p-values for overall survival for multiple TCGA cancer types using both CoCu8 and CoCu30 signatures. CESC - cervical squamous cell carcinoma; HNSCC - head and neck squamous cell carcinoma; LUSC – lung squamous cell carcinoma; PRAD - prostate adenocarcinoma; PAAD - pancreatic adenocarcinoma; LUAD - lung adenocarcinoma; KIRC - kidney clear cell carcinoma; COAD - colorectal adenocarcinoma; BLCA - bladder urothelial carcinoma; STES - esophagogastric carcinoma; SKCM – melanoma; BRCA - breast cancer. **B)** Kaplan-Meier overall survival analysis of HNSCC (left), CESC (center), LUSC (right) TCGA datasets stratified for CoCu30 first vs last quartile. Below each analysis are shown the corresponding numbers at risk, time in years. HNSCC HR=1.57 (95%CI 1.06-2.35), p-value=0.024. CESC HR=2.08 (95%CI 1.09-4.01), p-value=0.024. LUSC HR=1.78 (95%CI 1.23-2.59), p-value=0.0019. HR and CI were calculated using Cox regression. p-value was calculated using logRank test. **C)** Kaplan-Meier overall survival analysis of HNSCC HPV positive (left) and HNSCC HPV positive (right) TCGA datasets stratified for CoCu30 first vs last quartile. Below each analysis are shown the corresponding numbers at risk, time in years. HPV positive HR=5.47 (95%CI 1.76-17.0), p-value=0.0011. HPV negative HR=1.00 (95%CI 0.67-1.51), p-value=0.98. HR and CI were calculated using Cox regression. p-value was calculated using logRank test. **D)** Kaplan-Meier overall survival analysis of LUSC TRACERx dataset. Individual tumors stratified as high-, discordant or low-risk according to expression profile of CoCu30 signature across multiple regions, as previously described and stratified according to Biswas et al. ^60^. Below are shown the numbers at risk in years. HR=1.98 (95% CI 1.09-3.6), p-value=0.0152. HR and CI calculated using Cox regression and are referred to CoCu30 low vs CoCu30 high. p-value was calculated using logRank test.

**Figure S3:**
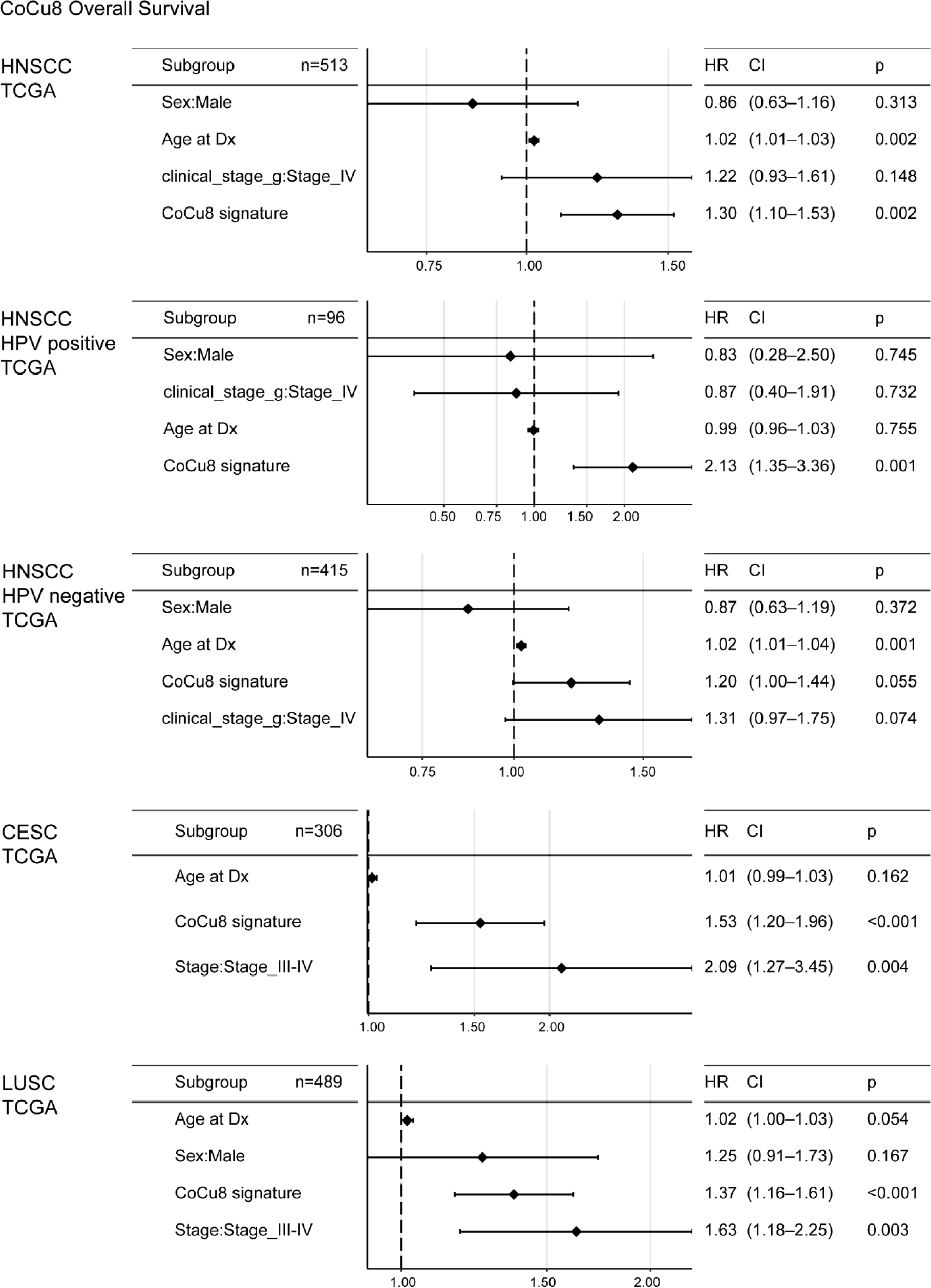
Multivariate analysis of CoCu8 overall survival. Forest plot showing Hazard Ratios, 95% confidence interval and p value calculated using multivariate Cox regression from patients with HNSCC (HPV positive and negative), LUSC and CESC from the TCGA cohort. Variables include: age (continuous, years), sex (male vs female, except for CESC as all patients were female), clinical stage (categorical) and the CoCu8 signature (continuous variable).

**Figure S4:**
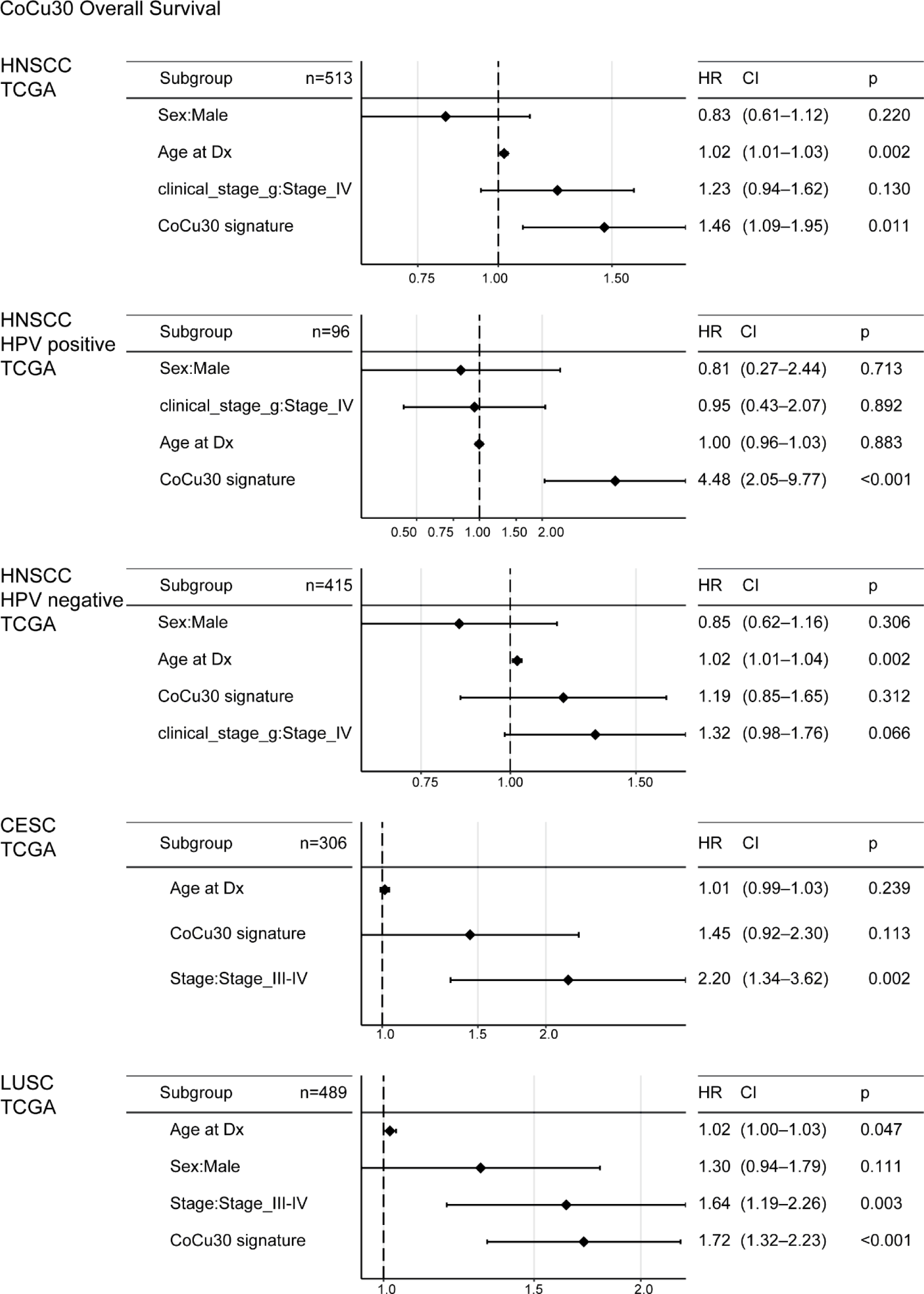
Multivariate analysis of CoCu30 overall survival. Forest plot showing Hazard Ratios, 95% confidence interval and p value calculated using multivariate Cox regression from patients with HNSCC (HPV positive and negative), LUSC and CESC from the TCGA cohort. Variables include: age (continuous, years), sex (male vs female, except for CESC as all patients were female), clinical stage (categorical) and the CoCu30 signature (continuous variable).

**Figure S5:**
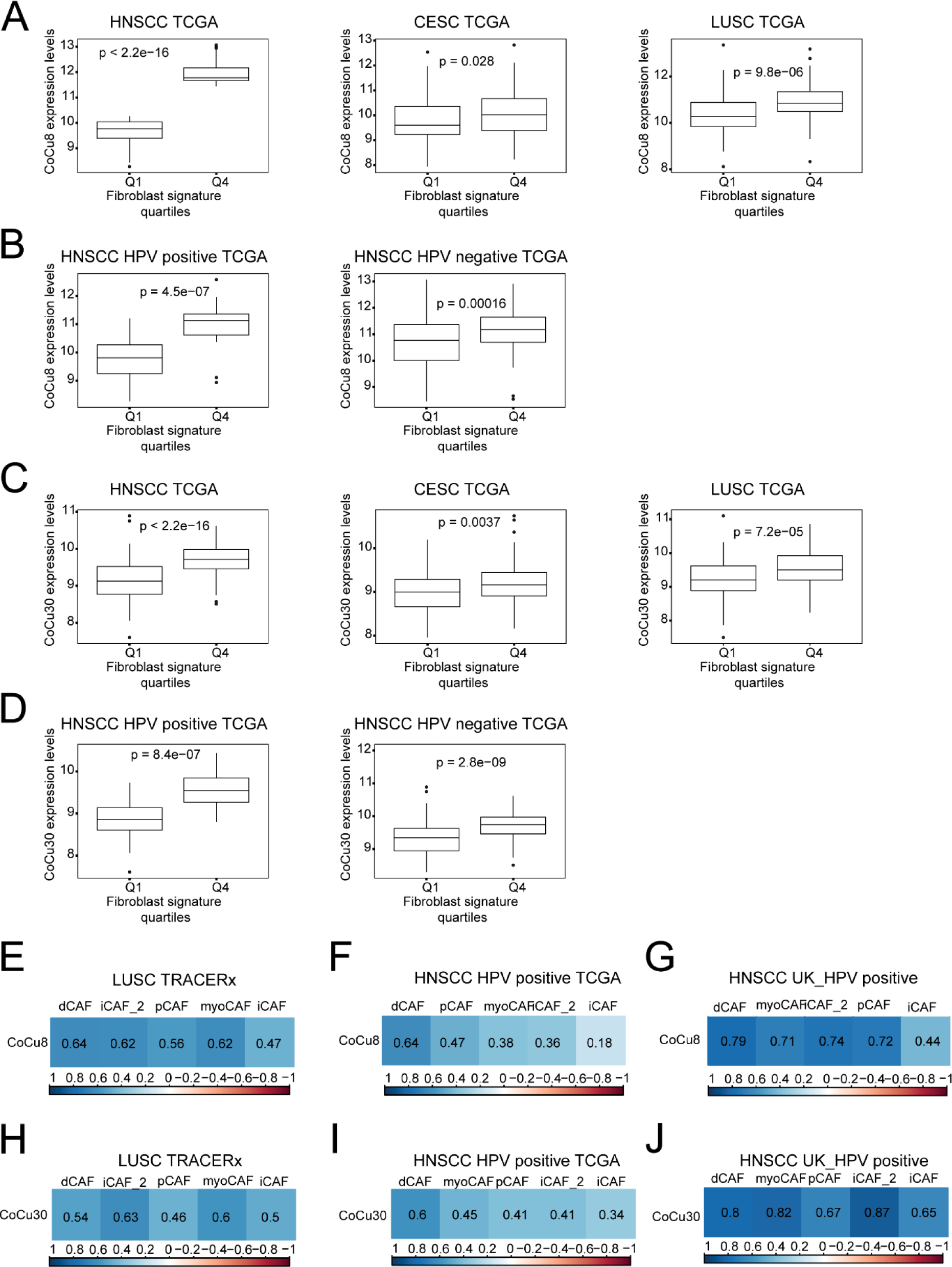
Fibroblast abundance correlates with CoCu8 / CoCu30 gene signature in different squamous cell carcinoma datasets. **A)** Box plot analysis of CoCu8 expression in HNSCC, CESC and LUSC TCGA separated by first and last quartile of fibroblast abundance via Methyl CIBERSORT deconvolution strategy. Independent Student’s t-test. **B)** Box plot analysis of CoCu8 expression in HPV positive and negative TCGA separated by first and last quartile of fibroblast abundance via Methyl CIBERSORT deconvolution strategy. Independent Student’s t-test. **C)** Box plot analysis of CoCu30 expression in HNSCC, CESC and LUSC TCGA separated by first and last quartile of fibroblast abundance via Methyl CIBERSORT deconvolution strategy. Independent Student’s t-test. **D)** Box plot analysis of CoCu30 expression in HPV positive and negative TCGA separated by first and last quartile of fibroblast abundance via Methyl CIBERSORT deconvolution strategy. Independent Student’s t-test. **E-J)** Correlation plot of different fibroblast subpopulations derived from Galbo et al. ^9^ with CoCu8 gene signature in LUSC TRACERx **(E)**, HPV positive HNSCC TCGA **(F)**, UK_HPV positive HNSCC **(G)** patients and with CoCu30 gene signature in LUSC TRACERx **(H)**, HPV positive HNSCC TCGA **(I)**, UK_HPV positive HNSCC **(J)** patients. The number inside the square represents the R, Spearman correlation coefficient. The color legend is shown at the bottom. All correlations are significant p-value<0.05.

**Figure S6:**
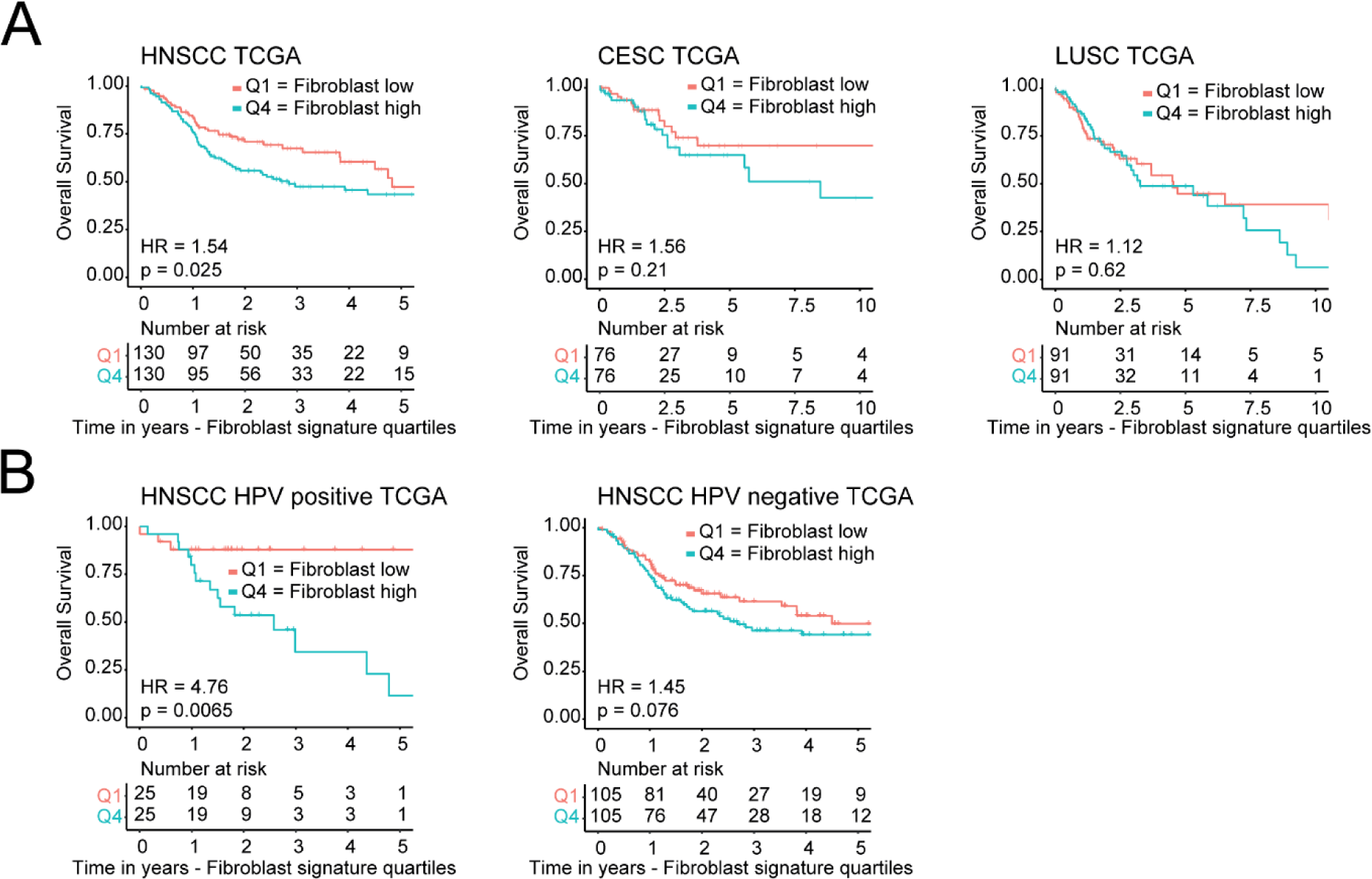
Fibroblast abundance meta-analysis on different squamous cell carcinoma datasets. **A)** Kaplan-Meier overall survival analysis of HNSCC (left), CESC (center), LUSC (right) TCGA datasets stratified for fibroblast abundance first vs last quartile. Below each analysis are shown the corresponding numbers at risk, time in years. HNSCC HR=1.54 (95%CI 1.05-2.25), p-value=0.025. CESC HR=1.56 (95%CI 0.77-3.17), p-value=0.21. LUSC HR=1.12 (95%CI 0.71-1.77), p-value=0.62. HR and CI were calculated using Cox regression. p-value was calculated using logRank test. **B)** Kaplan-Meier overall survival analysis of HNSCC HPV positive (left) and HNSCC HPV positive (right) TCGA datasets stratified for fibroblast abundance first vs last quartile. Below each analysis are shown the corresponding numbers at risk, time in years. HPV positive HR=4.76 (95%CI 1.36-16.5), p-value=0.0065. HPV negative HR=1.45 (95%CI 0.96-2.18), p-value=0.076. HR and CI were calculated using Cox regression. p-value was calculated using logRank test.

**Figure S7:**
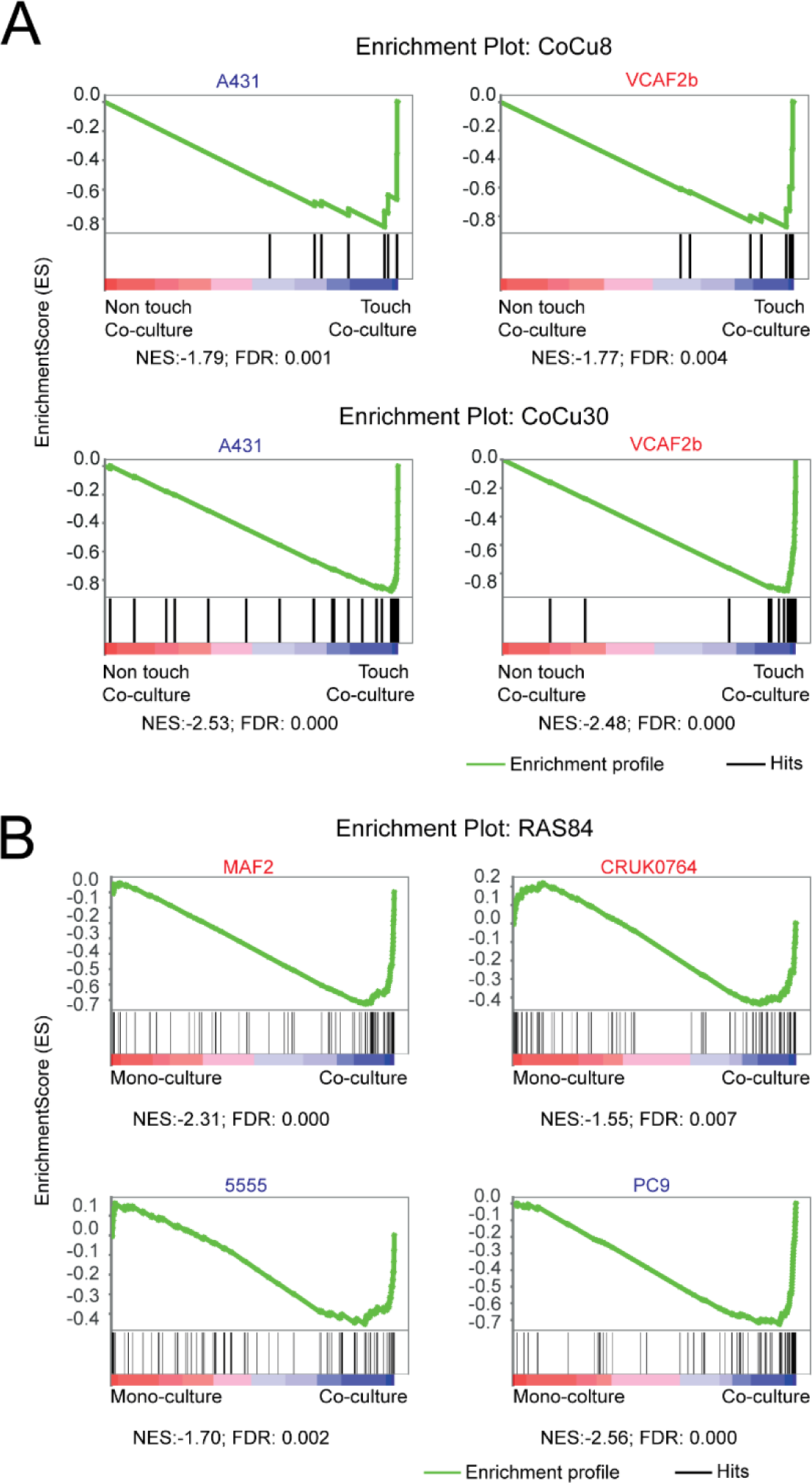
Enrichment of CoCu8, CoCu30 and RAS84 gene signatures in different co-culture conditions. **A)** Gene set enrichment analysis (GSEA) plot of CoCu8 gene signature (top) and CoCu30 (bottom) in co-culture indirect vs direct condition in A431 / VCAF2b transcriptomic dataset. NES and FDR are specified below each plot. **B)** Gene set enrichment analysis (GSEA) plot of RAS84 gene signature ^28^ in mono-culture and co-culture. NES and FDR are specified below each plot.

**Figure S8:**
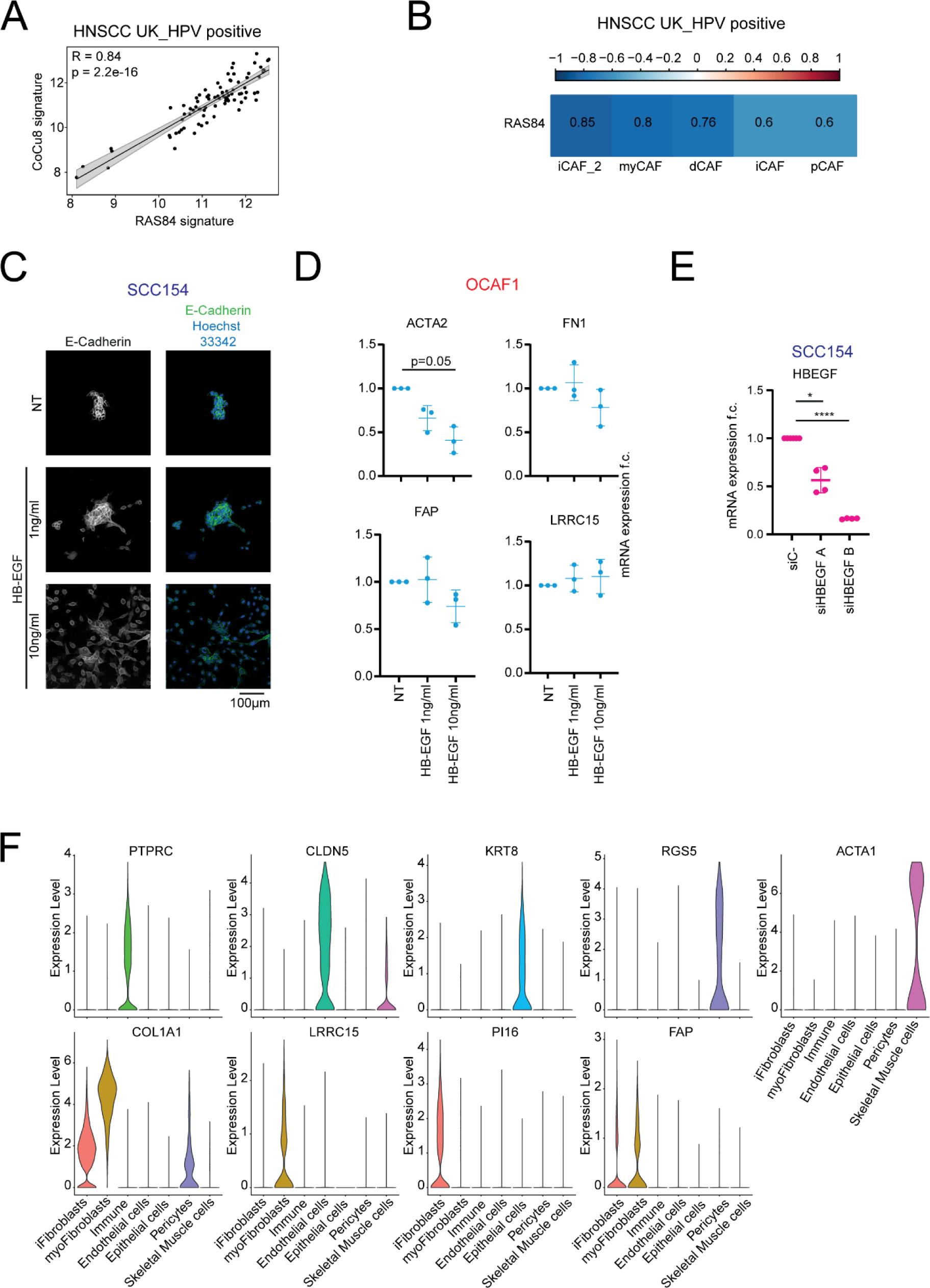
HB-EGF/RAS/MAPK activity in cancer cells and CAFs mono-culture vs co-culture. **A)** Correlation plot of CoCu8 and RAS84 expression levels in HNSCC HPV positive UK_HPV positive cohort. R is Spearman correlation coefficient. **B)** Correlation table of different fibroblast subpopulations derived from Galbo et al. ^9^ with RAS84 gene signature in the independent cohort UK_HPV positive of HNSCC HPV positive patients. The number inside the square represents the Spearman R, correlation coefficient. The color legend is shown on the top. All correlations are significant at p-value<0.001. n=97. **C)** Immunofluorescence staining of E-Cadherin and Hoechst 33342 for SCC154 mono-culture for the indicated treatments after 48h. **D)** qPCR analysis of *ACTA2*, *FAP*, *FN1* and *LRRC15* genes in OCAF1 mono-culture for the indicated treatments after 48h. mRNA expression is reported as mean ± standard deviation (SD) fold change difference over non-treated (NT) condition. Genes have been normalized over the average of *GAPDH*, *ACTB* and *RPLP0* housekeeping genes. n = 3 independent experiments. **E)** qPCR analysis of *HBEGF* gene in SCC154 after 96h of treatment with the indicated conditions. mRNA expression is reported as mean ± standard deviation (SD) fold change difference over siC-condition. Gene has been normalized over the average of *GAPDH*, *ACTB* and *RPLP0* housekeeping genes. n ≥ 4 independent experiments. Two tailed paired Student’s t-test. **F)** Violin plot of mRNA expression levels of the indicated genes for each cluster from Choi et al. ^33^.

**Figure S9:**
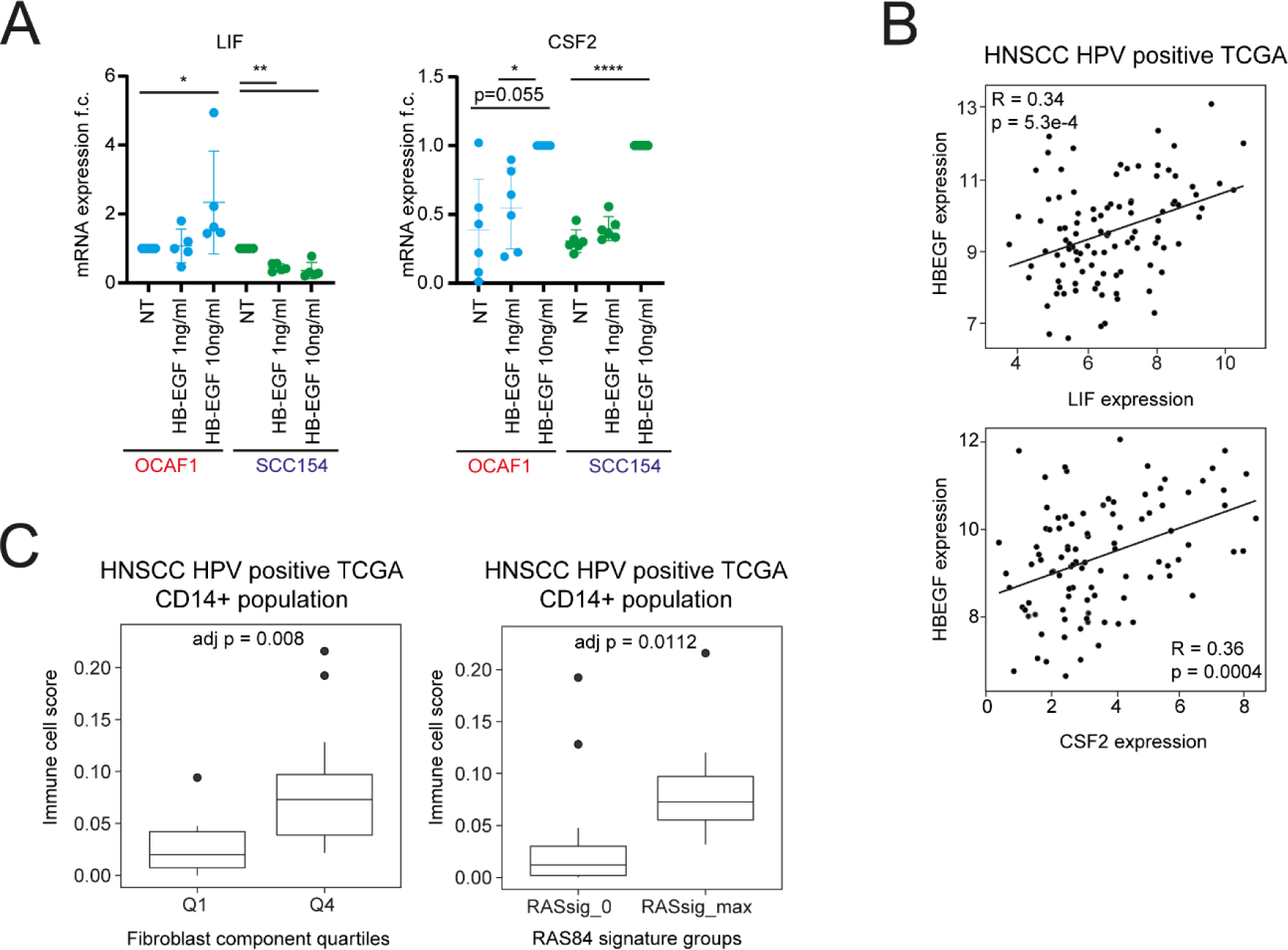
HB-EGF effect on cytokine production and monocyte/macrophage enrichment. **A)** qPCR analysis of *LIF* and *CSF2* genes in OCAF1 and SCC154 mono-cultures for the indicated treatments after 48h. mRNA expression is reported as mean ± standard deviation (SD) fold change difference over non-treated (NT) condition for each cell type with *LIF*, while for *CSF2* fold change difference is reported over Hb-EGF 10ng/ml treated sample. Genes have been normalized over the average of *GAPDH*, *ACTB* and *RPLP0* housekeeping genes. n ≥ 5 independent experiments. Two tailed paired Student’s t-test. **B)** Correlation plot of *HBEGF* with *LIF* (left) and *HBEGF* with *CSF2* (right) expression levels in HNSCC HPV positive TCGA dataset. R is Spearman correlation coefficient. **C)** (Right) Box plot analysis of immune cell absolute score via Methyl CIBERSORT deconvolution strategy in HNSCC HPV positive separated by first and last quartile of fibroblast abundance. Independent Student’s t-test. Bonferroni correction for multiple comparisons. (Left) Box plot analysis of immune cell absolute score via Methyl CIBERSORT deconvolution strategy in HNSCC HPV positive separated by RAS84_0 and RAS84_max. Student’s t-test Bonferroni correction for multiple comparisons.

**Figure S10:**
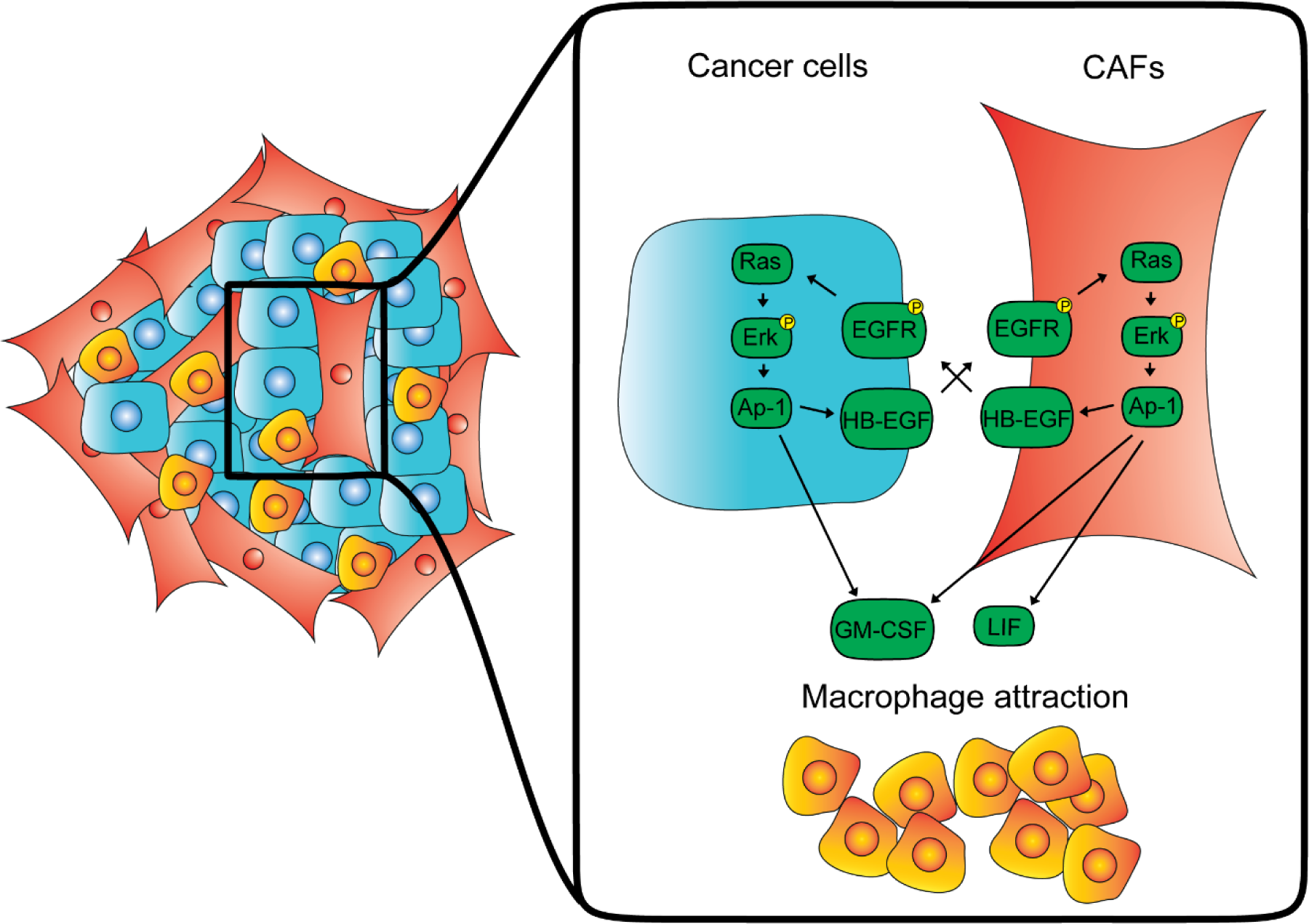
Schematic representation of the current cross-talk model of cancer cells – CAFs co-culture.

