## Supplementary Tables for "Cancer cell – fibroblast crosstalk via HB-EGF/EGFR/MEK signaling promotes macrophage recruitment in squamous cell carcinoma"

| Markers | Cell line: OCAF1 | Cell line: OCAF2 | Cell line: 796T |
|---|---|---|---|
| FGA | 19, 20 | 22, 26 | 19,24 |
| TPOX | 8, 11 | 8, 10 | 11,11 |
| D8S1179 | 13, 14 | 14, 15 | 11,12 |
| vWA | 14, 17 | 15, 17 | 17,17 |
| AMEL | X, Y | X, X | X,Y |
| Penta D | 13, 14 | 9, 13 | 10,12 |
| CSF1PO | 11, 12 | 11, 12 | 10,11 |
| D16S539 | 11, 13 | 11, 12 | 12,12 |
| D7S820 | 11, 12 | 10, 11 | 8,10 |
| D13S317 | 9, 13 | 11, 12 | 8,9 |
| D5S818 | 11, 11 | 12, 12 | 12,12 |
| Penta E | 12, 12 | 11, 18 | 8,13 |
| D18S51 | 14, 16 | 12, 16 | 11,15 |
| D21S11 | 28, 31 | 27, 28 | 29,31 |
| TH01 | 9.3, 9.3 | 6, 6 | 6,9.3 |
| D2S1358 | 15, 19 | 15, 15 | 15,15 |

**Supplementary table 1:** STR profile

| GENE | Species | Entrez Gene ID | Primer direction | Sequence (5' to 3') |
|---|---|---|---|---|
| JUNB | Homo sapiens | 3726 | Forward | AACAGCCCTTCTACCACGAC |
| JUNB | Homo sapiens | 3726 | Reverse | CAGGCTCGGTTTCAGGAGTT |
| FOS | Homo sapiens | 2353 | Forward | TGACTGATACACTCCAAGCGG |
| FOS | Homo sapiens | 2353 | Reverse | GGCAATCTCGGTCTGCAAAG |
| FOSB | Homo sapiens | 2354 | Forward | CCGTTGAATTGGAAACTGCT |
| FOSB | Homo sapiens | 2354 | Reverse | CAGAGGGAGAGAGACCATCG |
| CXCL2 | Homo sapiens | 2920 | Forward | CACAGTGTGTGGTCAACATTTC |
| CXCL2 | Homo sapiens | 2920 | Reverse | CACAGAGGGAAACACTGCAT |
| CXCL8 | Homo sapiens | 3576 | Forward | TCTGGCAACCCTAGTCTGCT |
| CXCL8 | Homo sapiens | 3576 | Reverse | AAACCAAGGCACAGTGGAAC |
| LIF | Homo sapiens | 3976 | Forward | CCTTTCCCTGGTCCCTACTC |
| LIF | Homo sapiens | 3976 | Reverse | CACCCGTAAGGCTTATTCCA |

| | | | | |
|---|---|---|---|---|
| CSF2 | Homo sapiens | 1437 | Forward | AAATGTTTGACCTCCAGGAGCC |
| CSF2 | Homo sapiens | 1437 | Reverse | ATCTGGGTTGCACAGGAAGTT |
| ACTB | Homo sapiens | 60 | Forward | AGAAAATCTGGCACCACACC |
| ACTB | Homo sapiens | 60 | Reverse | CAGAGGCGTACAGGGATAGC |
| GAPDH | Homo sapiens | 2597 | Forward | CAAAAGGGTCATCATCTCTGC |
| GAPDH | Homo sapiens | 2597 | Reverse | AGTTGTCATGGATGACCTTGG |
| RPLP0 | Homo sapiens | 6175 | Forward | AACCCAGCTCTGGAGAAACT |
| RPLP0 | Homo sapiens | 6175 | Reverse | CCCCTGGAGATTTTAGTGGT |
| ACTA2 | Homo sapiens | 59 | Forward | ACCCTGTTCCAGCCATCCTT |
| ACTA2 | Homo sapiens | 59 | Reverse | TGCCCCCTGATAGGACATTG |
| COL7A1 | Homo sapiens | 1294 | Forward | GGGCATCCAGCTACATCCTA |
| COL7A1 | Homo sapiens | 1294 | Reverse | GCTTGAGATCCCTGGAAGTG |
| FN1 | Homo sapiens | 2335 | Forward | CAAAGCAAGCCCGGTTGTTA |
| FN1 | Homo sapiens | 2335 | Reverse | CCCACTCGGTAAGTGTTCCC |
| LRRC15 | Homo sapiens | 131578 | Forward | AGGGACGGACACTGTACCTG |
| LRRC15 | Homo sapiens | 131578 | Reverse | GGGTACCATGGTGTTTCTGG |
| PDGFRA | Homo sapiens | 5156 | Forward | CTGGGTTTCCATCCTTGAG |
| PDGFRA | Homo sapiens | 5156 | Reverse | TAGTAGGCTTCCTGCGTGG |
| FAP | Homo sapiens | 2191 | Forward | TCAAAGAAGTATCCCTTGCTAATTCA |
| FAP | Homo sapiens | 2191 | Reverse | GCAATGACCATCCCTTCCTTAC |
| S100A4 | Homo sapiens | 6275 | Forward | AGGGACAACGAGGTGGACTT |
| S100A4 | Homo sapiens | 6275 | Reverse | CCCAACCACATCAGAGGAGT |

**Supplementary table 2:** primers

| PROTEIN | PRODUCER | CATALOG NUMBER | SPECIES | DILUTION |
|---|---|---|---|---|
| Phospho-p44/42 MAPK (Erk1/2) (Thr202/Tur204) | Cell Signaling | 4376 | Rabbit | 1:1000 in BSA 5% 1h at Room Temperature |

| | | | | |
|---|---|---|---|---|
| (20G11) | | | | |
| Erk1/2 (137F5) | Cell Signaling | 4695 | Rabbit | 1:1000 in milk 5% 1h at Room Temperature |
| Vinculin | Abcam | ab18058 | Mouse | 1:1000 in milk 5% 1h at Room Temperature |
| EGF Receptor (D38B1) XP | Cell Signaling | 4267 | Rabbit | 1:1000 in milk 5% 1h at Room Temperature |
| pEGFR Tyr1173 | Santa Cruz | SC-12351 | Goat | 1:1000 in BSA 5% O/N at 4oC |
| HB-EGF | R&D Systems | AF-259 NA | Goat | 1:500 in milk 5% O/N at 4oC |
| E-Cadherin | Cell Signaling | 3195 | Rabbit | 1:250 in BSA 3% O/N at 4oC |
| Rabbit anti-Goat IgG (H+L) Secondary Antibody, HRP | ThermoFisher | 31402 | Rabbit | 1:2000 in milk 5% 1h at Room Temperature |
| Goat anti-Mouse IgG (H+L) Secondary Antibody, HRP | ThermoFisher | 31430 | Goat | 1:2000 in milk 5% 1h at Room Temperature |
| Goat anti-Rabbit IgG (H+L) Secondary Antibody, HRP | ThermoFisher | 31460 | Goat | 1:2000 in milk 5% 1h at Room Temperature |
| Donkey anti-Rabbit IgG (H+L) Highly Cross-Adsorbed Secondary Antibody Alexa Fluor 555 | Invitrogen | A31572 | Donkey | 1:100 in BSA 3% 45 min at Room Temperature |

**Supplementary table 3:** antibodies
